# Estimation of Voxelwise Effective Connectivities: Applications to High Connectivity Sub-Regions within Hippocampal and within Corticostriatal Networks

**DOI:** 10.1101/039057

**Authors:** Ruben Sanchez-Romero, Joseph D. Ramsey, Jackson C. Liang, Kevin Jarbo, Clark Glymour

## Abstract

Standard BOLD connectivity analyses depend on aggregating the signals of individual voxel within regions of interest (ROIs). In certain cases, this aggregation implies a loss of valuable functional and anatomical information about sub-regions of voxels that drive the ROI level connectivity. We describe a data-driven statistical search method that identifies the voxels that are chiefly responsible for exchanging signals between regions of interest that are known to be effectively connected. We apply the method to high-resolution resting state functional magnetic resonance imaging (rs-fMRI) data from medial temporal lobe regions of interest of a single healthy individual measured repeated times over a year and a half. The method successfully recovered densely connected voxels within larger ROIs of entorhinal cortex and hippocampus subfields consistent with the well-known medial temporal lobe structural connectivity. To assess the performance of our method in more common scanning protocols we apply it to resting state fMRI data of corticostriatal regions of interest for 50 healthy individuals. The method recovered densely connected voxels within the caudate nucleus and the putamen in good qualitative agreement with structural connectivity measurements. We describe related methods for estimation of effective connections at the voxel level that merit investigation.

## Introduction

In one common method for estimating network connections from BOLD time series, voxels are clustered into regions of interest (ROIs) and averages or first principal components over voxels are used as variables. ROIs formed this way are chosen because, either on anatomical, experimental or statistical grounds, a cluster of voxels is thought to act coherently as a cause of coherent activity in other clusters, or to respond coherently as an effect of other clusters, or both. Connections are estimated at the ROI level rather than the voxel level because averaging voxel BOLD signals is thought to increase the signal-to-noise-ratio (Nieto-Castanon et al., 2003, Faria et al., 2012, Wong 2014). The result is logically peculiar: on the one hand, larger magnets and improvements in acquisition protocols are sought to increase fMRI spatial resolution; on the other hand, the BOLD signals from voxels are de-resolved into clusters from each of which a single variable is constructed.^1^ The result of this approach is a loss of potentially valuable information about the sub-regions of voxels within ROIs that are chiefly involved in ROI level effective connections. In what follows, we describe a procedure for resolving a given collection of ROIs into sub-sets of voxels that are chiefly responsible for ROI level effective connections. We illustrate the procedure with repeated single individual high-resolution resting state fMRI data from medial temporal lobe (MTL) cortices and the hippocampus, for which the relevant regions of interest, functions and effective connections are well studied (Amaral, 1993; Witter, 1993; Witter et al., 2000; Zeineh et al., 2000, 2001, 2012; Libby et al., 2012; Yushkevich et al., 2009, 2010; Ekstrom et al., 2009; Malykhin et al., 2010; Preston et al., 2010; Yassa et al., 2010); and with 50 individuals resting state fMRI data from a corticostriatal network confirmed by tractography on diffusion spectrum imaging data (Jarbo and Verstynen, 2015).

## Voxelwise Conditional Independence

Our procedure considers the setting in which a ROI level causal graph has been previously established, whether by experimental intervention on brain tissues or by statistical analysis of brain imaging signals, or by other means, but it is not known which voxels in distinct ROIs are directly connected with one another. We assume the voxels driving the connectivity between two ROIs are those voxels in each region with the largest number of direct functional connections to voxels in the other. Accordingly, we need a method to count voxelwise *direct* connections between effectively connected ROIs.

BOLD signals of two individual voxels, *x* and *y*, respectively within two effectively connected regions of interest R_x_ and R_y_, may be correlated for any of several reasons. Physiological activity in one voxel may directly affect activity in another; or there may be a chain of voxels that are causally intermediate between x and y, or there may be one or more common causes of activity in x and y by other voxels.

Here we are interested in using a given ROI level causal structure to find the voxels within a ROI that are directly and actively signaling other voxels in a ROI directly effectively connected to the first, as in figure 1, while avoiding spurious associations that result from unaccounted common causes or intermediate variables in a causal chain.

**Figure 1.**
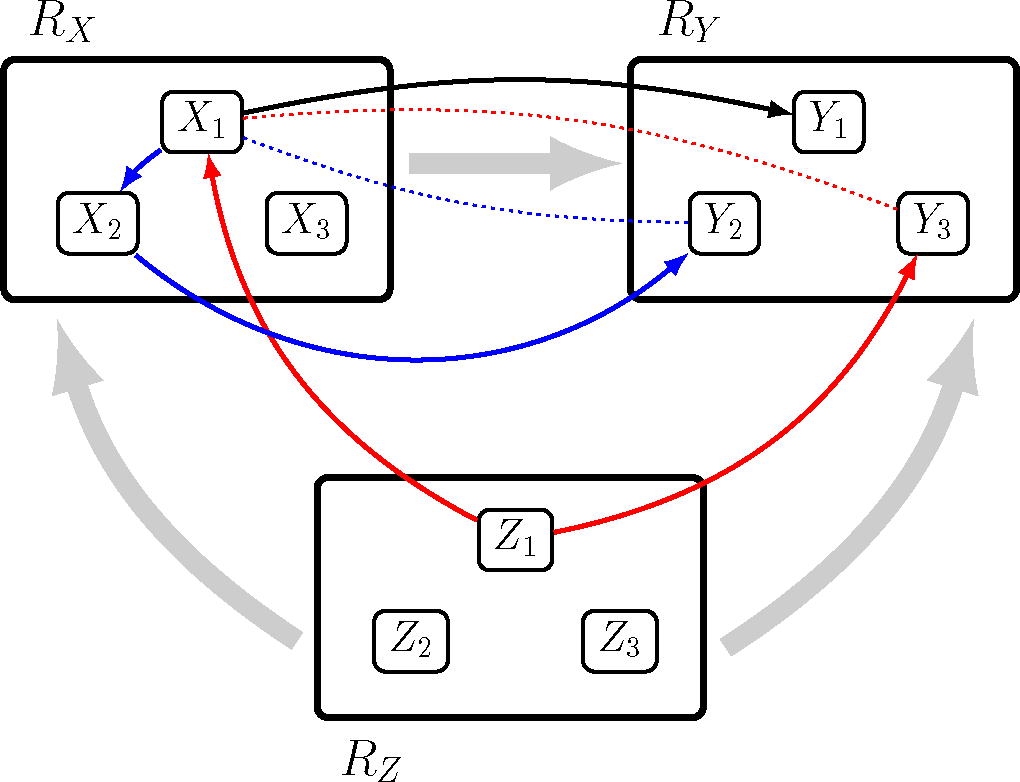
Causal graph at ROI level and voxel level. Thick grey arrows represent the ROI level effective connectivity where region of interest Rz is a common cause of R_x_ and R_y_, and R_x_ is a *direct cause* of Ry. Voxel X1 in R_x_ is a direct cause of voxel Y1 in R_y_ (black arrow). Voxel Z1 in Rz is a *common* cause of voxels X1 and Y3 in R_x_ and R_y_ respectively (red arrows). Red dotted line represents the spurious association between voxel X1 and voxel Y3 produced by the common cause Z1. The causal chain going from voxel X1 to voxel X2 in R_x_ to voxel Y2 in R_y_ is represented by blue arrows. The blue dotted line represents the spurious association between voxels X1 and Y2 produced by the causal chain X1 to X2 to Y2. To estimate the direct associations of voxel X1 with all voxels in R_y_ we need to condition on the common cause Z1 in Rz and on the intermediate voxel in the causal chain, X2 in R_x_, to avoid spurious associations, X1-Y3 and X1-Y2.

For each pair of voxels *x, y*, respectively in ROIs R_x_, R_y_ we estimate their independence or dependence conditional on all other voxels in R_x_ and R_y_ and all other voxels in ROIs that directly effectively connect both R_x_ and R_y_, except for ROIs that are common effects of R_x_ and R_y_.^2^ For each voxel *x* in R_x_ we count the number of voxels in R_y_ found to be conditionally dependent on *x.* We refer to this number as the *connectivity degree* for *x* in the R_x_ - R_y_ pair. Similarly, we obtain the *connectivity degree* for *y* in the R_x_ - R_y_ pair. The voxels in any pair of effectively connected regions of interest may be compared simply by their connectivity degree or most highly connected subsets may be characterized. To select the subset of voxels with the highest number of direct functional connections, the connectivity degree of all the voxels in R_x_ and R_y_ are respectively partitioned into 2 sets using k-means clustering. One set corresponds to highly connected voxels and the other set to voxels with lower or no connections in the R_x_ - R_y_ pair. Voxels in the highly connected set yield the *R_x_ - R_y_ high communication sub-region* in R_x_ and in R_y_ respectively. If in the ROI level effective connectivity, R_x_ → R_y_, a high communication sub-region in R_x_ can be thought as a massive sender of information to R_y_, while a high communication subregion in R_y_ is a massive receiver of information from Rx. Besides the bipartite division, we describe the voxelwise connectivity degree across ROIs.

A region of interest, R_x_, may have distinct high communication sub-regions for other regions of interest with which R_x_ is directly connected according to the given causal graph, or it may have overlapping communication sub-regions which suggest areas where afferent and/or efferent information may be interacting. The voxels not in any high communication sub-region may serve as relays between high communication sub-regions or may be voxels that were misclustered in forming the region of interest—we do not propose to determine which is the case.

## Algorithm for the Voxelwise Conditional Independence

The conditional independence test we use is based on inverting the covariance matrix over a set of variables to obtain an estimate of the partial correlation between two variables conditional on all the other variables in the set. However, if the number of variables exceeds the number of datapoints, which is typically the case for a single scanning session, the inversion of the covariance matrix is not possible due to rank deficiency. When there are sufficient scans of the same individual or of multiple individuals correctly aligned, under the same experimental conditions, the multiple scan data can be concatenated to guarantee that the number of datapoints exceeds the number of variables. We follow this approach and concatenate the necessary scans to satisfy the dimensionality condition before inputting the data in the Voxelwise Conditional Independence algorithm.

Let **R** be a set of non-overlapping ROIs <R_1_,…,R_*t*_>, and let G be a directed graph of effective connections between members of **R.** We denote the region of interest containing voxel *x* by R_x_. A *trek* t between two nodes, R_x_, R_y_ of G is any directed path between R_x_, R_y_, or any pair of directed paths, one terminating in R_x_ and the other in R_y_ such that the pair of paths intersect in one and only one node. Any node on a trek t between R_x_, R_y_ is said to *trek separate* R_x_, R_y_ with respect to t. If R_x_, R_y_ are adjacent in G, let <R_z1_,…,R_z*n*_> be any minimal subset of **R** that separates *all* treks between R_x_, R_y_ except for the direct connection between R_x_, R_y_.

We apply a conditional independence test to each pair of voxels, *x, y*, respectively in two directly effectively connected regions of interest, R_x_, R_y_, conditional on all other voxels in R_x_, R_y_ and all voxels on the minimal subset of regions of interest that trek separate R_x_, R_y_ (except for the trek that is the direct connection between R_x_, R_y_.)

#### Voxelwise Conditional Independence Algorithm

For a pair of regions of interest R_x_ and R_y_ and conditioning separating subset <R_z1_,…,R_z*n*_>: (R_x_, R_y_ | R_z1_,…,R_z*n*_)

1. Compute the covariance matrix C over the voxels in R_x_, R_y_, R_z1_,…,R_z*n*_.
2. Invert C yielding precision matrix Q.
3. For each voxel *x* in R_x_, *y* in R_y_, calculate the partial correlation: r(*x, y*) = -Q(*x, y*) / sqrt(Q(*x, x*)Q(*y, y*)).
4. For each voxel *x* in R_x_, *y* in R_y_, calculate the Fisher Z transform of r(*x, y*): *f*(*x, y*) = sqrt(N - 1 - dimension(Q))*(ln(1+r(*x, y*)) - ln(1 - r(*x, y*))), where N is the number of datapoints.
5. For each voxel *x* in R_x_, *y* in R_y_, calculate the p-value of f(*x, y*): p(*x, y*) = 2*(1 - CDF(f(*x, y*)), where CDF is the cumulative distribution function for the standard Gaussian distribution *N*(0, 1).
6. Make a list P of all the p-values calculated in step 5.
7. Sort P from low to high and apply False Discovery Rate at an α level.
8. For each voxel *x* in R_x_, *y* in R_y_, compute cd(*x*, R_y_):=number of voxels in R_y_ judge not independent of voxel *x* from step 7; cd(*y*, R_x_):= number of voxels in R_x_ judge not independent of voxel *y* from step 7.

We supplement the Voxelwise Conditional Independence algorithm with a classification into “ high communication sub-regions” using the connectivity degree cd(.) computed in step 8:

9. For regions of interest, R_x_, R_y_, apply k-means clustering with k=2 to the cd(*x*, Ry) of all voxels *x* in R_x_ and respectively to the cd(*y*, R_x_) of all voxels *y* in R_y_.
10. Define the k-means cluster with the higher cd(*x*, R_y_) values as the R_x_ - R_y_ high communication sub-region in R_x_ (respectively for R_y_).

The algorithm can obviously be stopped at step 8 to provide the distribution of connectivity degrees of cross-ROI connections without continuing the somewhat arbitrary bipartite division of voxels in step 9 and 10. Instead voxelwise connectivity degrees can be shown as spatial heat maps, as we do for illustrative cases in figures 5 – 9. Some technical remarks about the algorithm are given in the appendix.

## Application to resting state medial temporal lobe data

The functional segregation of the medial temporal lobe (MTL) has been extensively studied, although ambiguities remain. Figure 2 shows a simple diagram of functional components and their main structural connections, based on Squire et al., (2004) and Preston and Wagner (2007).

**Figure 2.**
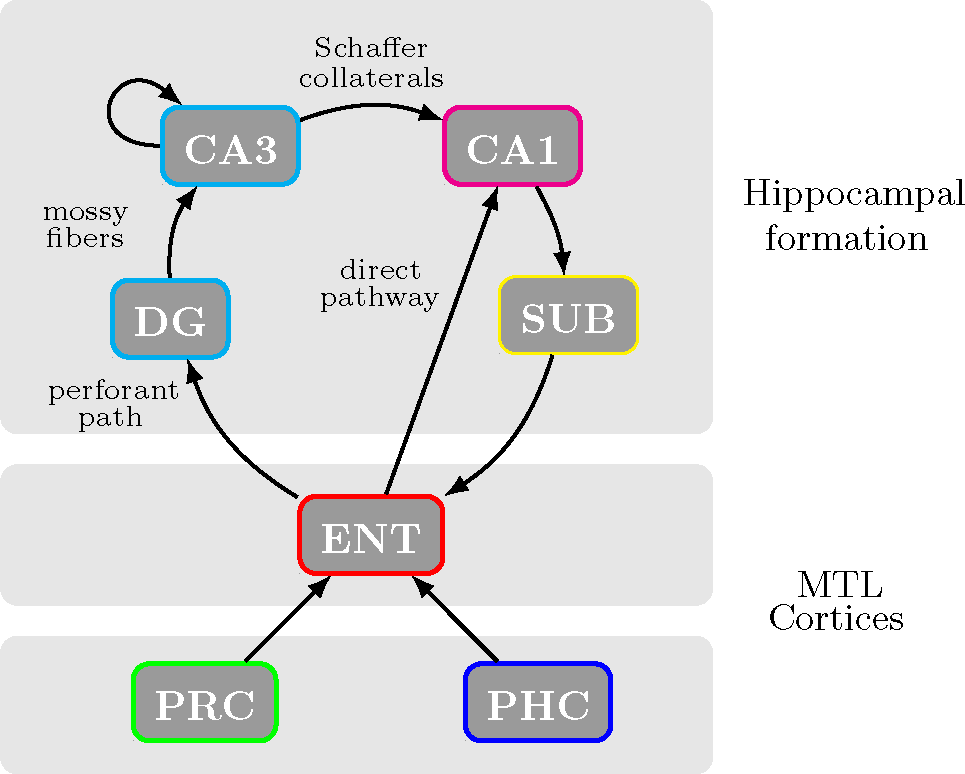
One interpretation of ground truth for the medial temporal lobe effective connectivity, including MTL cortices (perirhinal (PRC), parahippocampal (PHC) and entorhinal (ENT) cortices) and hippocampal formation (dentate gyrus (DG), CA3, CA1 and subiculum (SUB)).

There are however, finer partitions of the components of the medial temporal lobe that reveal some of the difficulties of functional connectivity analyses in this region of the brain. As an example, consider the entorhinal cortex, as shown in figure 3. This diagram suggests that if a ROI is created measuring only layers IV and V of the entorhinal cortex (EC in the diagram) the significant direct association with the dentate gyrus (DG) will not be found, or alternatively, measuring only layers I-III, no direct association with the subiculum (SUB) will be found.

**Figure 3.**
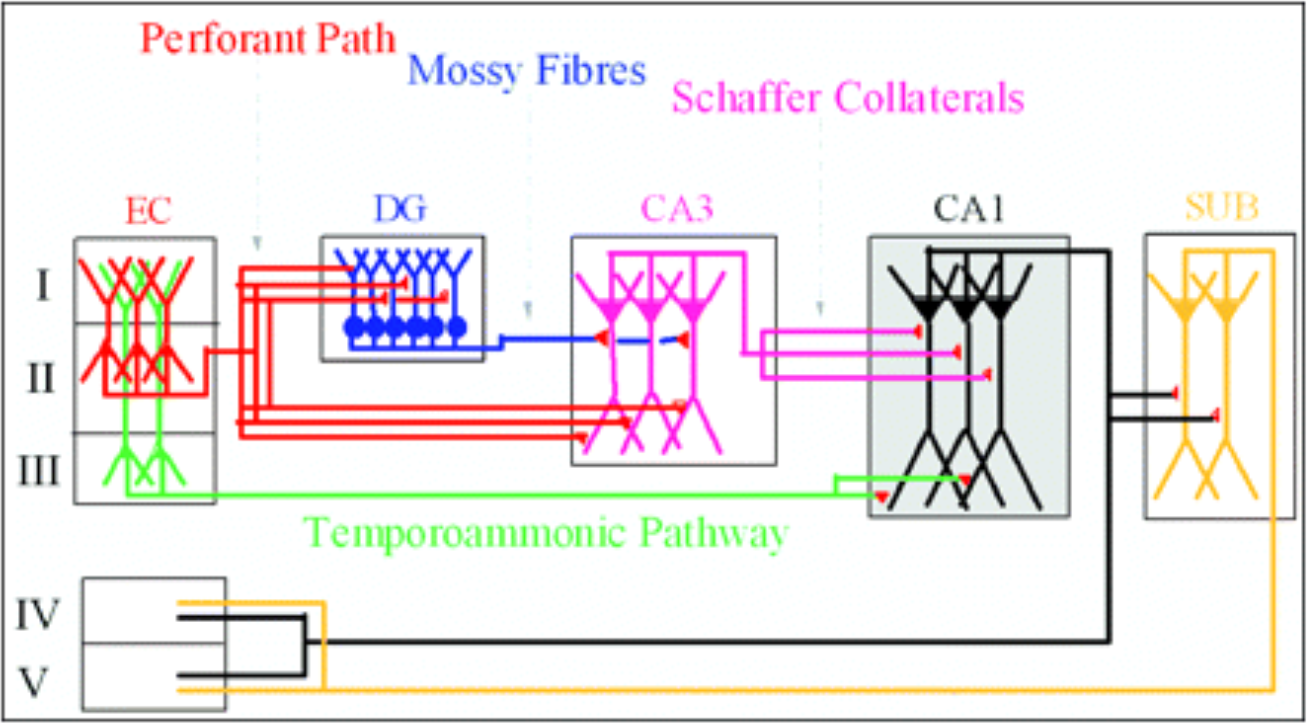
Schematic diagram of entorhinal/hippocampal synaptic pathways. Layers of entorhinal cortex (EC) are shown. Layers I-III have pathways with DG, CA3 and CA1. Layers IV-V have pathways with CA1 and subiculum. Taken from Coulter et al., (2011), © 2011 The Authors. Journal compilation, © 2011 The Physiological Society; with permission from John Wiley & Sons, Inc.

Obtaining proper ROIs for the medial temporal lobe, in particular for the hippocampus subfields, is an ongoing challenge (Yushkevich et al., 2015). Limits on the spatial resolution of functional imaging and the anatomical complexity of the hippocampus make difficult to obtain functional ROIs that correspond exactly to the anatomy of the subfields. The dentate gyrus, for example, is hardly separable in functional imaging from CA3 and CA2, resulting in a comprehensive ROI labeled CA32DG (Zeineh et al., 2000; Ekstrom et al., 2009; Preston et al., 2010). In addition, other studies have evidenced the complexity of the medial temporal lobe circuitry, such as, differences in functional connectivity profiles of anterior and posterior areas of CA1, CA32DG and subiculum with parahippocampal and perirhinal cortices (Libby et al., 2012); and connectivity differences of medial and lateral entorhinal cortex with cortical inputs and hippocampus subfields (Kerr et al., 2007; Canto et al., 2012).

The data used here were acquired, preprocessed, and provided to us by Russell Poldrack as part of the MyConnectome Project (myconnectome.org). We provide general information about the acquisition and preprocessing. Full details are available in Laumann et al. (2015) and Poldrack et al. (2015). MRI data were obtained repeatedly from one healthy individual over the course of 18 months. Scanning was performed in a fixed schedule, subject to availability of the participant. Scans were performed at fixed times of day; Mondays at 5 pm, and Tuesdays and Thursdays were performed at 7:30 am. Imaging was performed on a Siemens Skyra 3T MRI scanner using a 32- channel head coil. T1- and T2-weighted anatomical images were acquired using a protocol patterned after the Human Connectome Project (Van Essen et al., 2012). Anatomical data were collected on 14 sessions through 4/30/2013, with a one-year follow up collected on 11/4/2013. T1-weighted data were collected using an MP-RAGE sequence (sagittal, 256 slices, 0.8 mm isotropic resolution, TE=2.14 ms, TR=2400 ms, TI=1000 ms, flip angle = 8 degrees, PAT=2, 7:40 min scan time). T2-weighted data were collected using a T2-SPACE sequence (sagittal, 256 slices, 0.8 mm isotropic resolution, TE=565 ms, TR=3200 ms, PAT=2, 8:24 min scan time). Resting state fMRI was performed using a multiband EPI (MBEPI) sequence (Moeller et al., 2010) (TE = 30 ms, TR=1160 ms, flip angle = 63 degrees, voxel size = 2.4 mm x 2.4 mm x 2 mm, distance factor=20%, 68 slices, oriented 30 degrees back from AC/PC, 96 x 96 matrix, 230 mm FOV, MB factor=4, 10 min scan length). Starting with session 27 (12/3/2012), the number of slices was changed to 64 because of an update to the multiband sequence that increased the minimum TR beyond 1160 ms for 68 slices. A total of 104 resting state fMRI scanning sessions were acquired; 12 were pilot sessions using a different protocol, and additional 8 were excluded based on poor signal, leaving a total of 84 usable sessions. Functional data were preprocessed including intensity normalization, motion correction, atlas transformation, distortion correction using a mean field map, and resampling to 2mm atlas space. No spatial smoothing was applied.

Regions of interest in the medial temporal lobe were defined manually according to procedures established by the Preston Laboratory at The University of Texas at Austin (Liang et al., 2013). ROIs were defined bilaterally for subiculum, CA1, CA32DG, entorhinal cortex, perirhinal cortex and parahippocampal cortex. The high resolution T2 anatomical images of the MyConnectome Project allowed a more reliable delineation of hippocampal subfields in the body, head and tail of the hippocampus; Insausti and Amaral (2004) and Duvernoy et al., (2005) were used as anatomical guidelines. In addition, adjustments to the standard delineation of perirhinal cortex were made following Ding and Van Hoesen (2010). Figure 4 shows lateral and medial views of the 3D rendered left hemisphere ROIs. Right hemisphere ROIs are shown in figure S1 in supplementary material.

**Figure 4.**
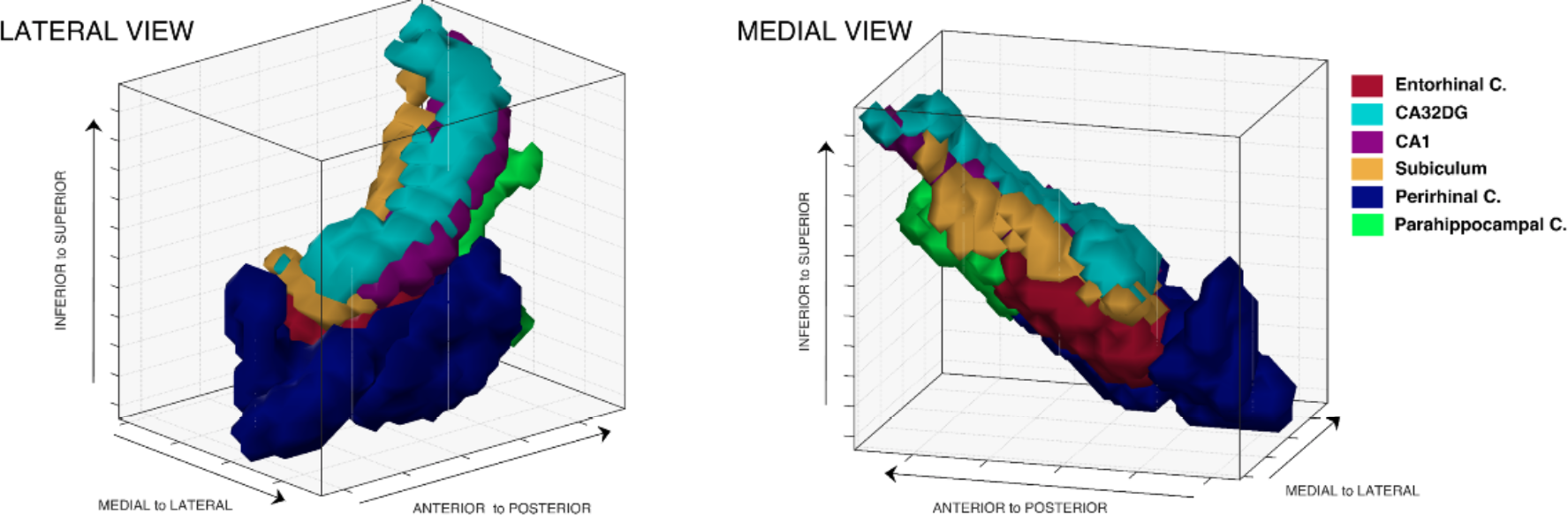
Lateral and medial views of 3D rendered left hemisphere medial temporal lobe showing ROIs for MTL cortices and hippocampus subfields.

Following the directed graph in figure 2, we ran the Voxelwise Conditional Independence algorithm for the following five pairs of dependence relations conditional on their corresponding separating subset: (ENT, CA32DG | SUB), (CA32DG, CA1 | ENT), (CA1, SUB | ENT), (SUB, ENT | CA1), (ENT, CA1 | SUB, CA32DG). To explore the robustness of the algorithm results, we created four new datasets by concatenating the first ten scanning sessions, concatenating the last ten scanning sessions, concatenating ten randomly chosen sessions, and concatenating the first five sessions. Results were highly similar in all four datasets for both the left and right hemisphere data. (Figure S2 - S8 in supplementary material.)

Using the first ten scanning sessions, the resulting high communication sub-regions are illustrated in figures 5 - 9 for the left hemisphere for each of the five pairs of dependent ROIs mentioned above. Each figure shows a 3D voxel space representation of the corresponding pair of ROIs, heatmaps of the voxelwise connectivity degree and the resulting high communication sub-regions. For visualization purposes we present exploded views by translating the position of one of the two ROIs across the medial-lateral and/or inferior-superior axes. See movies 1- 5 for rotating exploded views of high communication sub-regions.

For the medial temporal lobe data we observe high communication sub-regions with tight spatial coherency and consistent location in neighboring areas of directly connected regions of interest. We note, as expected, that voxels in a given high communication sub-region have direct cross-ROI connections with voxels in the corresponding high communication sub-region but we also observe a number of cross-ROI direct connections with voxels throughout the body of the ROI. (See figure S12 in supplementary material for a summary of distances between connected voxels.) Our high communication sub-regions are in excellent agreement with the high resolution diffusion tensor imaging results reported in Zeineh et al., (2012). In particular, the high communication sub-regions coincide with areas that show high density of fiber tracts for the corresponding medial temporal lobe pathways. (See figures 6-2, 6-4, 6-5 and 6-6 in Zeineh et al., (2012).)

**Figure 5.**
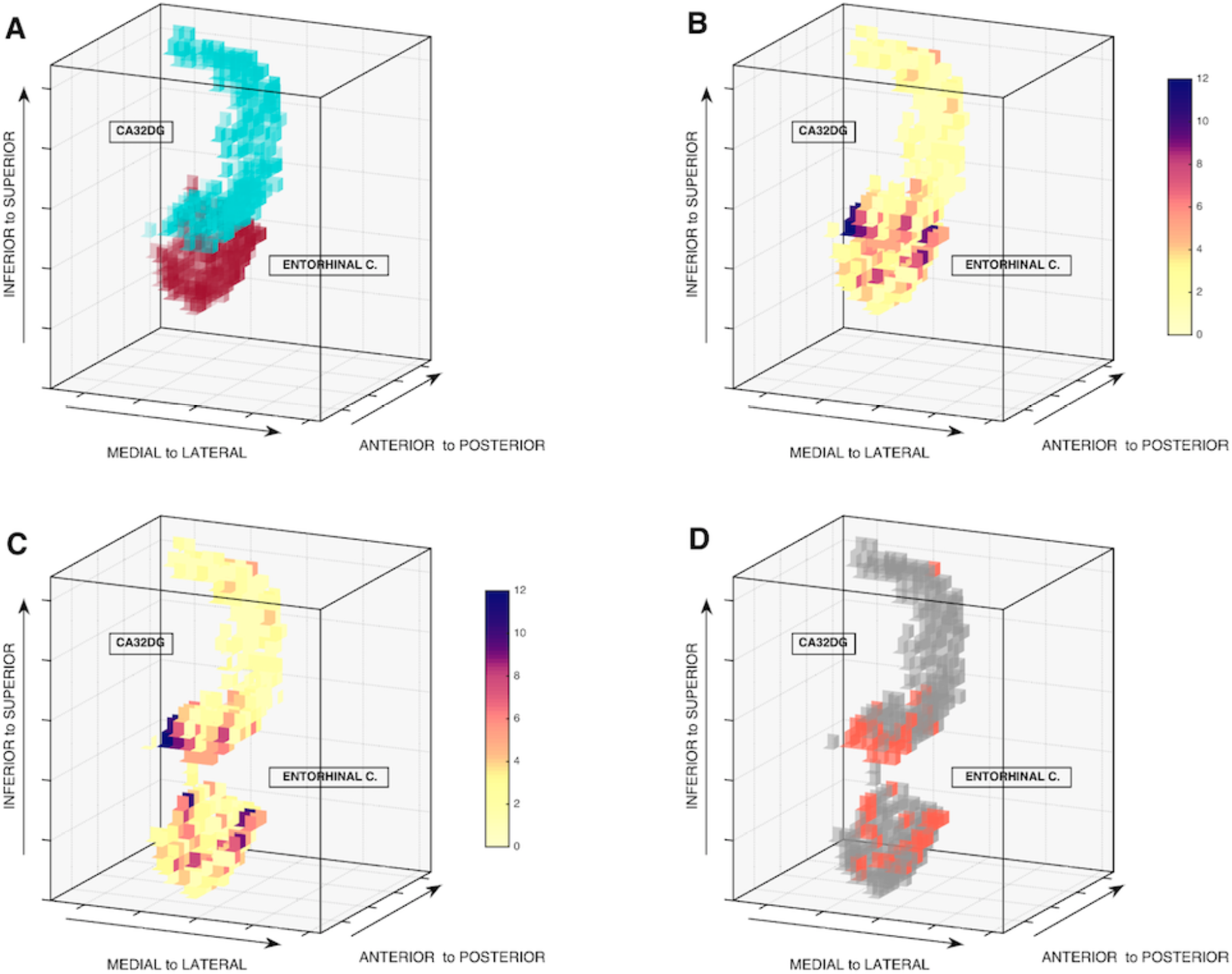
Left hemisphere 3D voxel space representation of (A) entorhinal cortex and CA32DG regions of interest; (B) voxelwise connectivity degree heatmaps for each region of interest (darker colors imply higher degree); (C) exploded view of connectivity degree heatmaps; (D) exploded view of high communication sub-regions (orange). Communication sub-regions are located in the inferior anterior portion of CA32DG and in the outmost medial and lateral sections of the entorhinal cortex. See movie 1.

**Figure 6.**
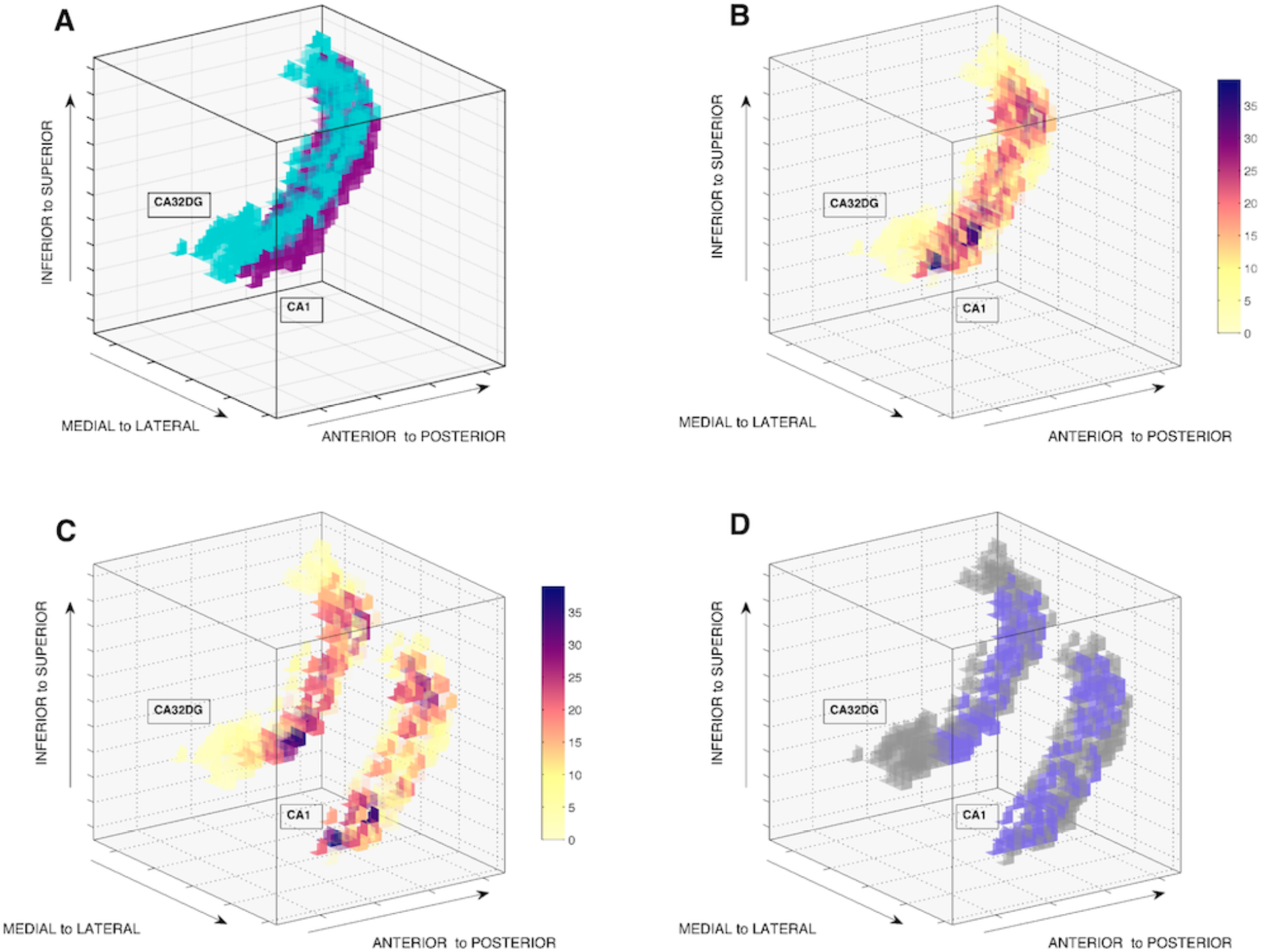
Left hemisphere 3D voxel space representation of (A) CA32DG and CA1 regions of interest; (B) voxelwise connectivity degree heatmaps for each region of interest (darker colors imply higher degree); (C) exploded view of connectivity degree heatmaps; (D) exploded view of high communication sub-regions (purple). The communication sub-regions are remarkably large in both regions of interest. In CA1 it includes a big stripe from superior to inferior areas. Conversely, in CA32DG it runs from superior to inferior areas, ending where the CA1 region ends. See movie 2.

**Figure 7.**
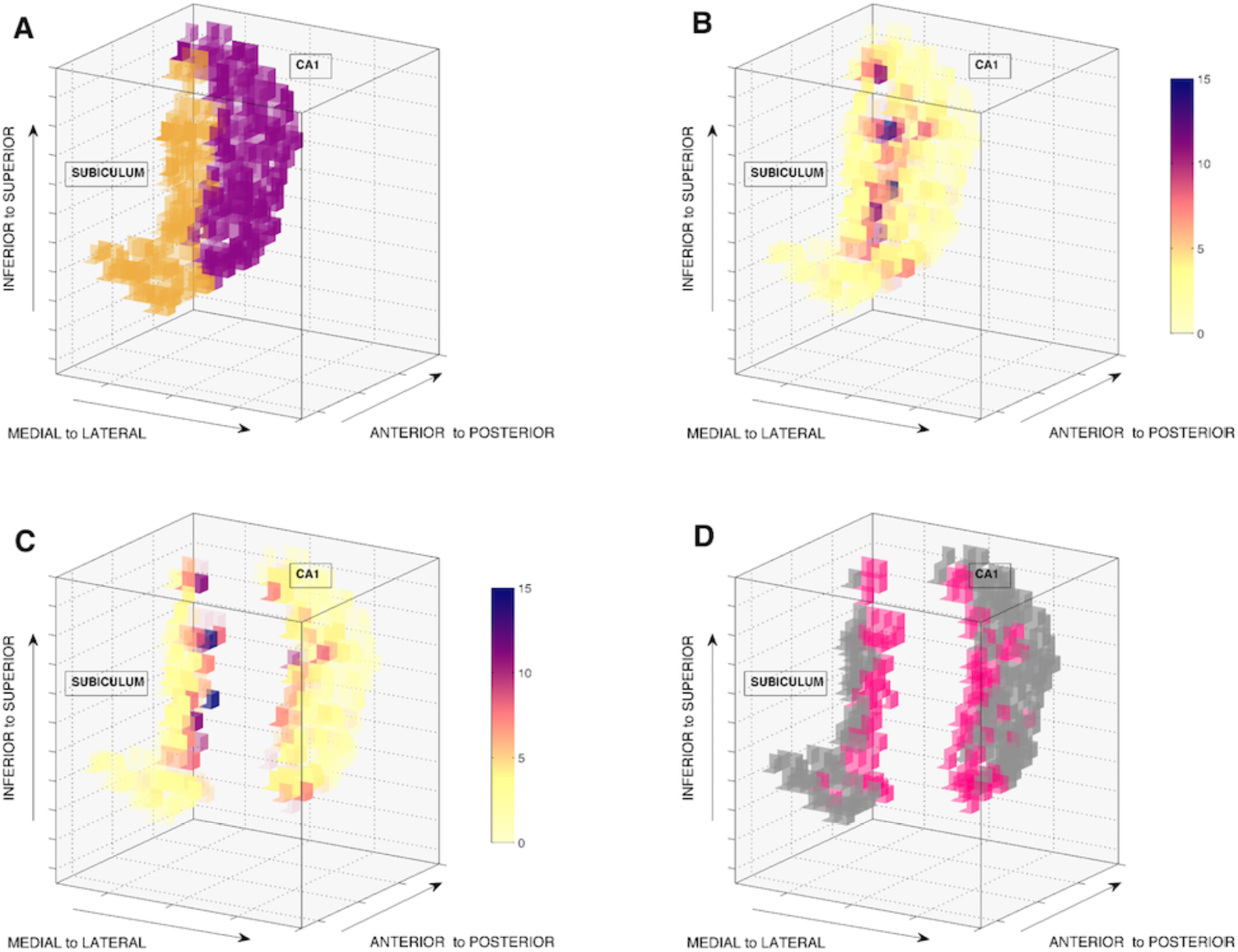
Left hemisphere 3D voxel space representation of (A) CA1 and subiculum regions of interest; (B) voxelwise connectivity degree heatmaps for each region of interest (darker colors imply higher degree); (C) exploded view of connectivity degree heatmaps; (D) exploded view of high communication sub-regions (pink). In the subiculum the communication sub-region includes inferior to top superior lateral areas. Conversely in CA1 the sub-region encompasses from inferior to top superior medial areas. See movie 3.

**Figure 8.**
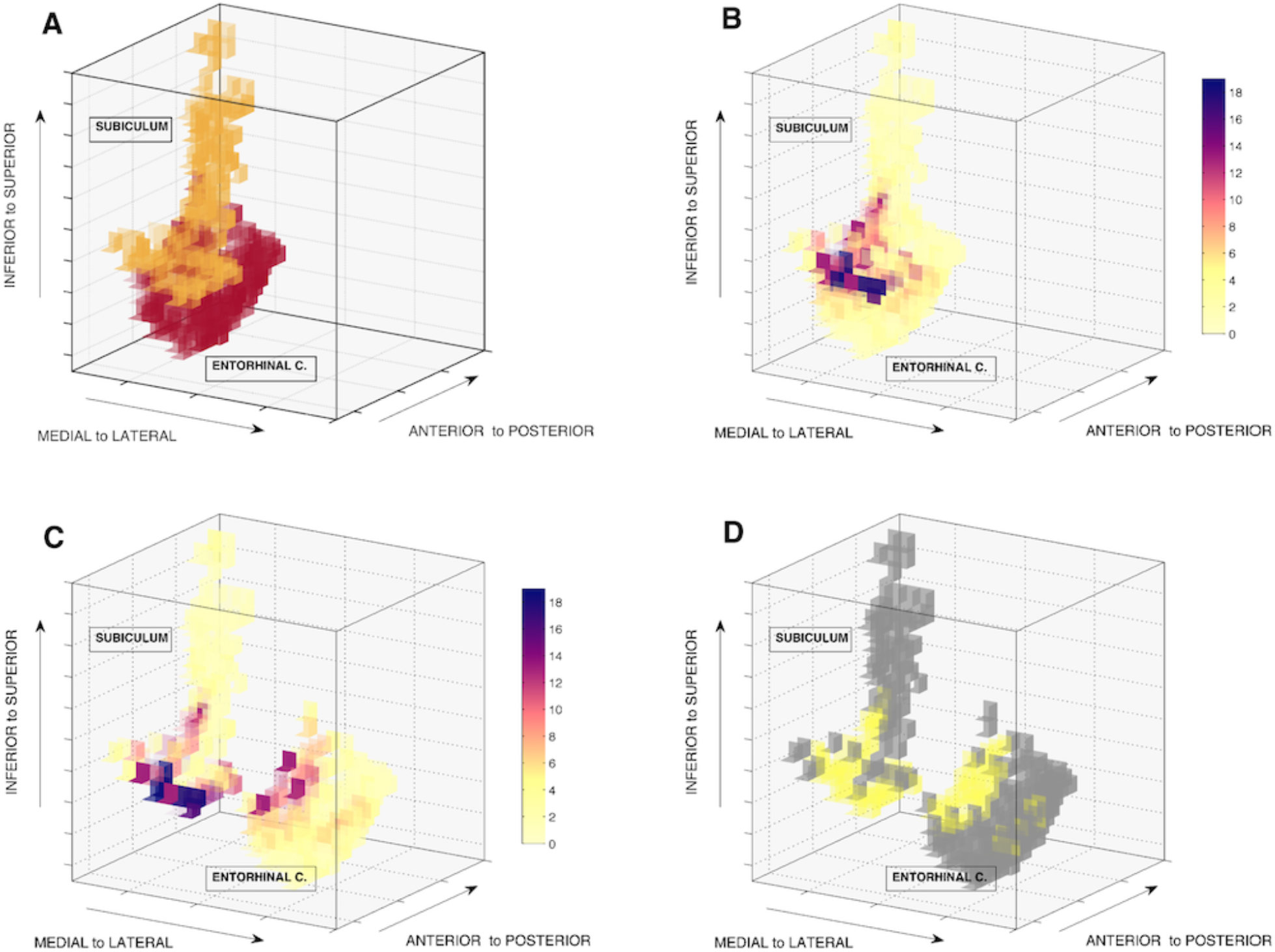
Left hemisphere 3D voxel space representation of (A) subiculum and entorhinal cortex regions of interest; (B) voxelwise connectivity degree heatmaps for each region of interest (darker colors imply higher degree); (C) exploded view of connectivity degree heatmaps; (D) exploded view of high communication sub-regions (yellow). In the entorhinal cortex the communication sub-region is located mainly in the medial superior part and more sparsely in the lateral part. In the subiculum it is located in its inferior lateral and medial part. See movie 4.

**Figure 9.**
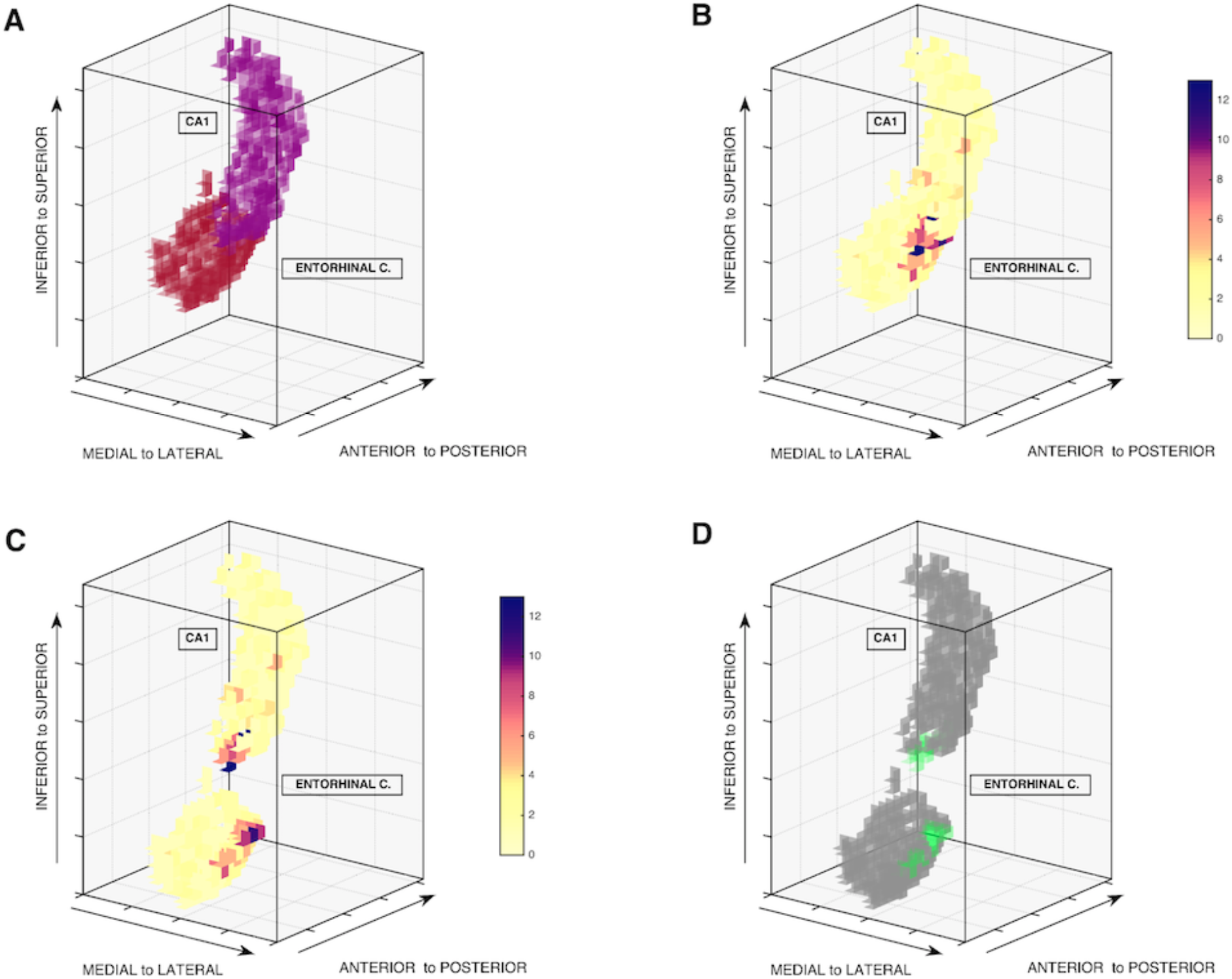
Left hemisphere 3D voxel space representation of (A) entorhinal cortex and CA1 regions of interest; (B) voxelwise connectivity degree heatmaps for each region of interest (darker colors imply higher degree); (C) exploded view of connectivity degree heatmaps; (D) exploded view of high communication sub-regions (green). The communication sub-regions are very compact in both ROIs. In entorhinal cortex it encompasses the utmost posterior lateral area and in CA1 the utmost inferior area. See movie 5.

High communication sub-regions in a particular region of interest may overlap indicating voxels with direct functional connectivity with more than one ROI. To illustrate this case we show in figure 10 the region of interest for left hemisphere entorhinal cortex and its high communication sub-regions with subiculum, CA1 and CA32DG, indicating the entorhinal voxels where overlapping occurs.

**Figure 10.**
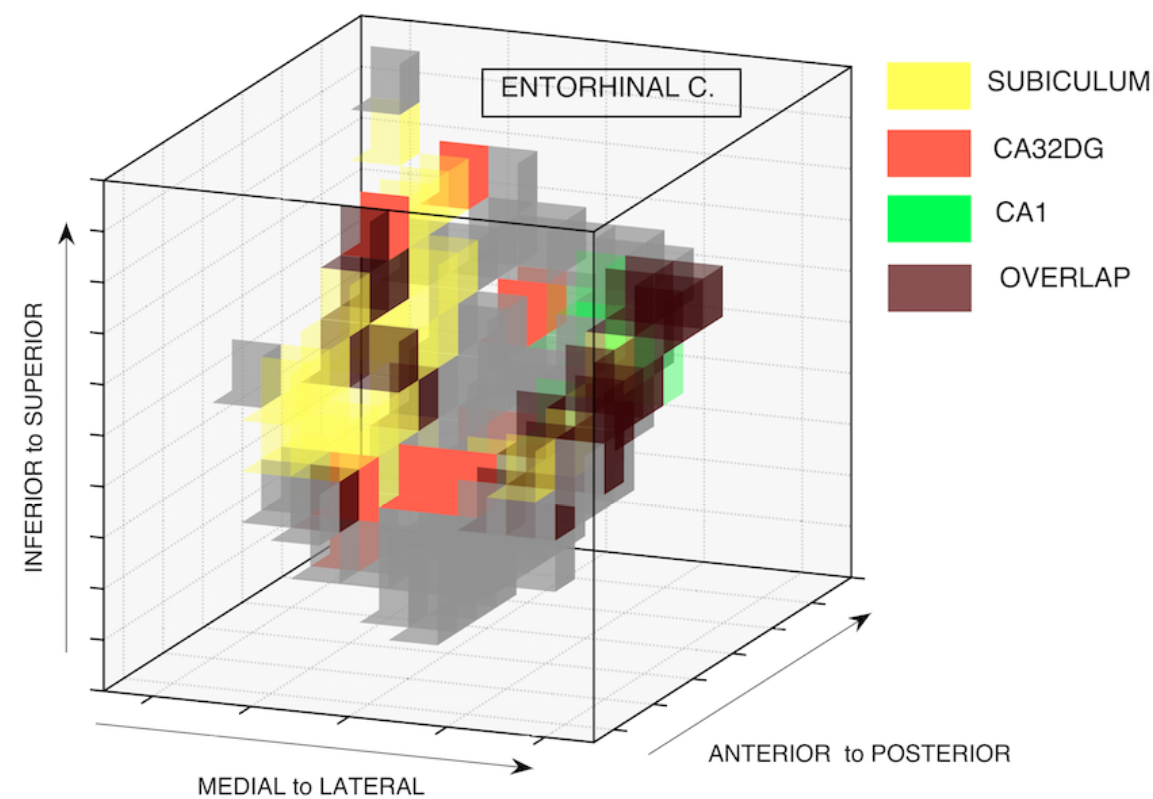
Left hemisphere 3D voxel space representation of entorhinal cortex, showing communication sub-regions with subiculum (yellow), CA1 (green) and CA32DG (orange), and voxels where overlapping of sub-regions occur (brown). Sub-regions with subiculum and CA32DG overlap in medial areas of the entorhinal cortex. Sub-regions with CA1 and CA32DG overlap in lateral areas.

## Sensitivity to registration issues

With multiple scans, whether of the same individual or different individuals, the procedure we have described raises questions of the accuracy of co-registration when the data from multiple scanning sessions are concatenated, as we did here. In particular, if non-equivalent voxels from various concatenated scans are incorrectly co-registered to the same standard space coordinates, estimation of the influence of a voxel in one ROI on voxels in another ROI will not come from concatenated signals of correctly equivalent voxels but from a mixture of signals of non-equivalent voxels. This problem is relevant given that mixture of signals has been reported as one of the most detrimental conditions for inference of fMRI functional and effective connectivity (Smith et al., 2011).

To test the sensitivity of the procedure to such effects, we simulate two cases of incorrect co-registration in the single individual medial temporal lobe data. In the first case, for each of the first ten concatenated sessions, we randomly shifted each voxel in each ROI by zero or one voxel step in each of the x, y, and z coordinates. For the second case, the same process was carried out but shifting each voxel by zero, one, or two voxel steps in each coordinate. In both cases, the resulting estimated high communication sub-regions were qualitatively identical to those obtained with the original data, but with slightly reduced connectivity degree for each voxel. We illustrate these results in figure 11, where for the left hemisphere entorhinal cortex we show the high communication sub-region with subiculum for the original data (zero shifting) and the second case data (zero, one or two voxel shifting). In both cases the sub-regions are qualitatively similar in location and density. We also show in figure 11 connectivity degree heatmaps for entorhinal cortex voxels. As previously mentioned, in the second case data (zero, one or two voxel shifting) the voxels in the communication sub-region have smaller connectivity degree than in the original data (zero shifting).

**Figure 11.**
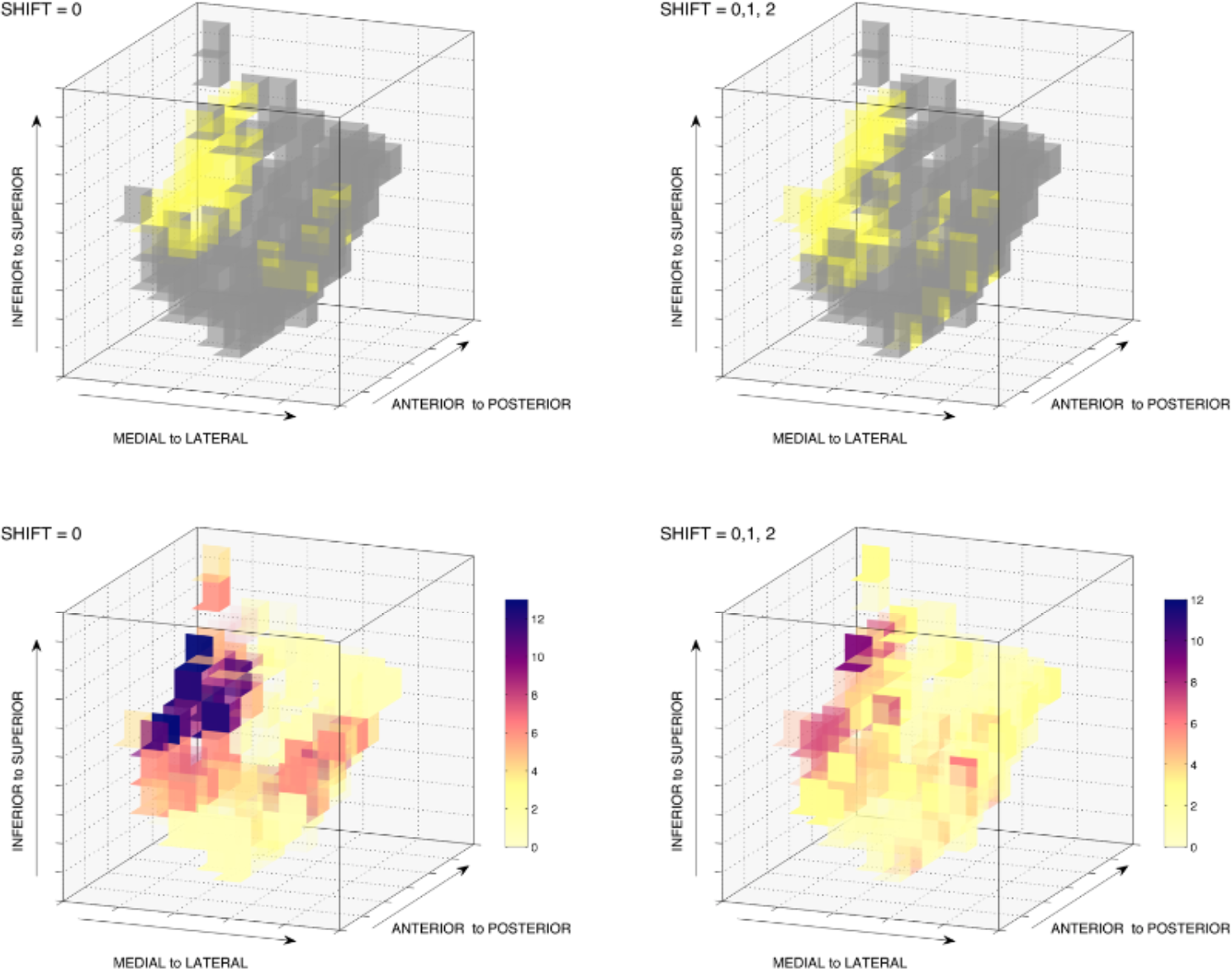
Left hemisphere 3D voxel space representation of entorhinal cortex showing, (top row) high communication sub-region with subiculum for original data (shift = 0) and second case data (shift = 0,1,2); (bottom row) connectivity degree heatmaps for original data (shift = 0) and second case data (shift = 0,1,2). The high communication sub-region is spatially similar in both cases but with differences in connectivity degrees.

## Application to resting state corticostriatal data

Using deterministic fiber tractography on diffusion spectrum imaging data for 59 individuals, Jarbo and Verstynen (2015) describe the intrahemispheric structural connectivity of inputs to the caudate nucleus and putamen from the orbitofrontal (OFC), dorsolateral prefrontal (DLPFC) and posterior parietal cortices. They show that anatomical connections coming from OFC, DLPFC and posterior parietal cortex converge in respective regions of the caudate and the putamen, suggesting the presence of a network that integrates reward, executive control and spatial attention during spatial reinforcement learning.

The Jarbo and Verstynen (2015) structural connectivity network together with well-established knowledge about the direction of the flow of information in corticostriatal pathways (Wilson, 1989) can be represented with the graphical structure in figure 12.

**Figure 12.**
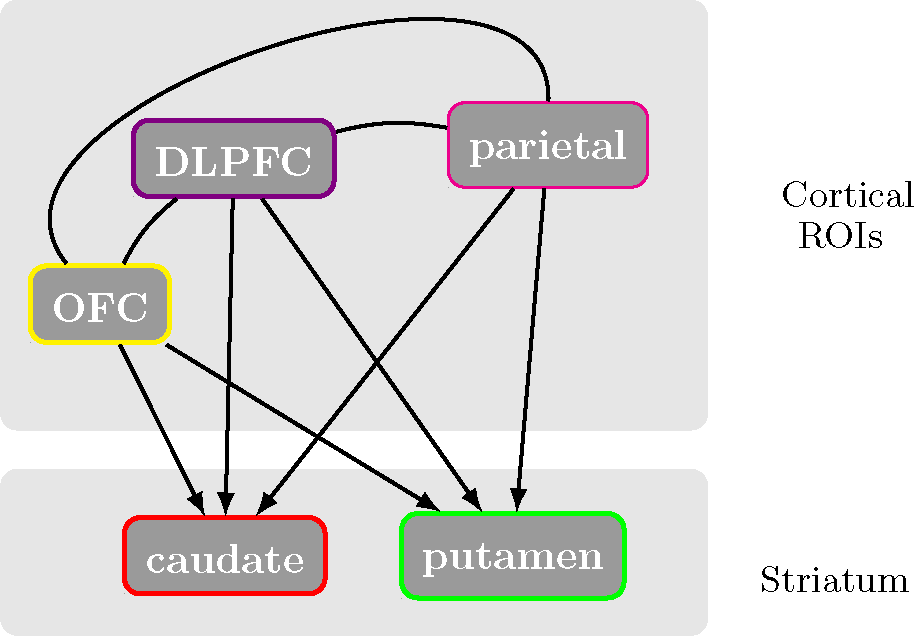
Graphical structure for selected regions of interest in a corticostriatal circuitry. The cortical ROIs, OFC, DLPFC and parietal are direct causes of the caudate and the putamen. Cortical ROIs are interconnected but we do not specify the direction of the interaction.

In accordance with the graphical structure in figure 12, we run the Voxelwise Conditional Independence algorithm for three conditional independence relations for the caudate nucleus, (caudate, OFC | DLPFC, parietal), (caudate, DLPFC | OFC, parietal), and (caudate, parietal | OFC, DLPFC); and three for the putamen (putamen, OFC | DLPFC, parietal), (putamen, DLPFC | OFC, parietal) and (putamen, parietal | OFC, DLPFC). We concatenate resting state fMRI data for 50 individuals^3^ described in Jarbo and Verstynen (2015).

Anatomical scans and raw resting state fMRI scans were provided to us by the Cognitive Axon Lab in Carnegie Mellon University. We include here general information about acquisition. Full details are available in Jarbo and Verstynen (2015). Anatomical T1-weighted high-resolution images acquired with an MPRAGE sequence (1mm isotropic voxels, 176 slices). Resting state functional data consisted of 210 T2*-weighted volumes with a BOLD contrast with echo planar imaging (EPI) sequence (TR= 2000ms, TE = 29ms, voxel size =3.5mm isotropic, field of view = 224 x 224 mm, flip angle = 79°).

We preprocessed the resting state functional data using FEAT (FMRI Expert Analysis Tool) Version 6.00, part of FSL (FMRIB’s Software Library, www.fmrib.ox.ac.uk/fsl): motion correction using MCFLIRT (Jenkinson et al., 2002); non-brain removal using BET (Smith, 2002); grand-mean intensity normalization of the entire 4D dataset by a single multiplicative factor; highpass temporal filtering (Gaussian-weighted least-squares straight line fitting, with sigma=50.0s) and no spatial smoothing to prevent voxelwise BOLD signal mixing. We apply the artifact removal procedure FIX (Salimi-Korshidi et al., 2014; Griffanti et al., 2014) using ICA components obtained with MELODIC 3.14 (Beckmann and Smith, 2004), conservative threshold level of 5, and the standard training dataset, as suggested in the user guide (fsl.fmrib.ox.ac.uk/fsl/fslwiki/FIX/UserGuide). The cleaned functional data was registered to FSL’s MNI152 standard space (2mm isotropic voxels) using linear registration FLIRT (Jenkinson and Smith, 2001; Jenkinson et al., 2002) with spline interpolation.

In contrast to Jarbo and Verstynen (2015) who used the SRI24 space multichannel atlas (Rohlfing et al., 2010) to select ROIs for their diffusion imaging analysis, we used the AAL atlas (Tzourio-Mazoyer et al., 2002) defined in the MNI space of the functional data. The six cortical ROIs (three per hemisphere) were created as indicated in Jarbo and Verstynen (2015): OFC was formed as an aggregation of gyrus rectus (Rectus); ventromedial prefrontal cortex (Frontal_Med_Orb); and opercular, orbital, and triangular parts of the inferior frontal gyrus (Frontal_Inf_Oper, Frontal_Inf_Orb, Frontal_Inf_Tri). DLPFC was formed as an aggregation of dorsal and orbital middle and superior frontal gyri (Frontal_Mid, Frontal_Mid_Orb, Frontal_Sup, Frontal_Sup_Orb). Parietal cortex as an aggregation of superior and inferior parietal lobules (Parietal_Sup, Parietal_Inf); angular gyrus (Angular) and supramarginal gyrus (SupraMarginal). Caudate and putamen ROIs were used as included in the AAL atlas. Figure 13 shows 3D rendered left and right hemisphere AAL ROIs for caudate and putamen.

**Figure 13.**
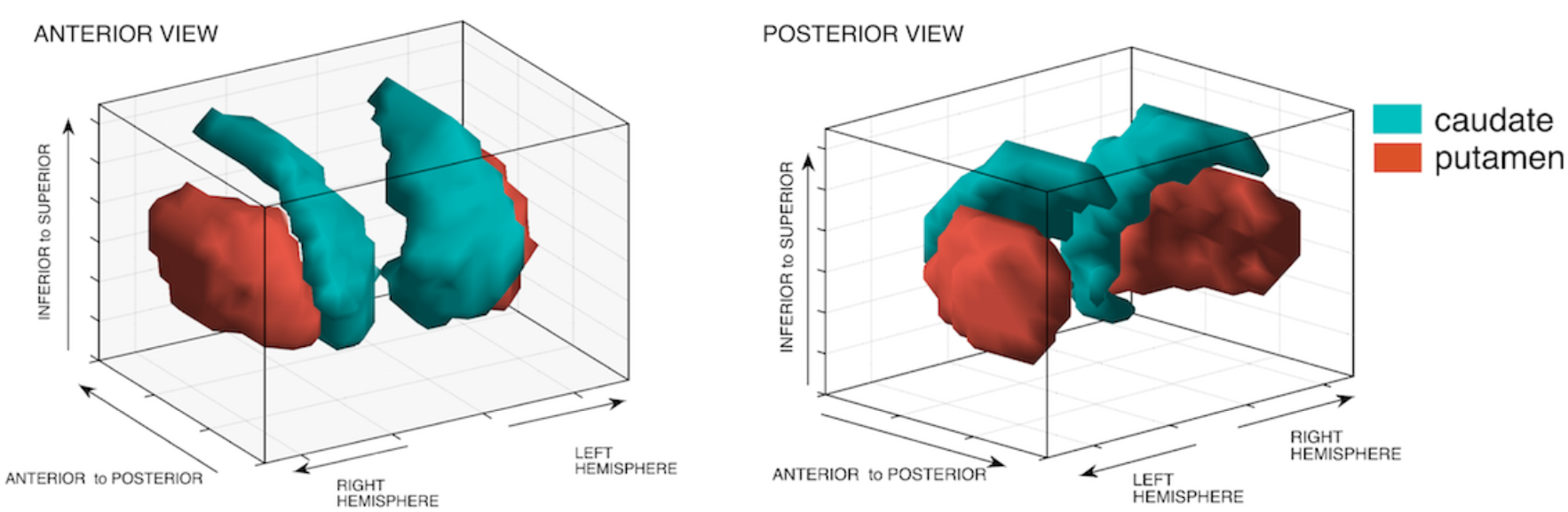
Anterior and posterior views of 3D rendered left and right hemisphere caudate and putamen AAL ROIs.

Given the number of voxels contained in each ROI in MNI space (2mm isotropic voxels), the covariance matrices to estimate the voxelwise conditional independence relations contain approximately 25,000 variables. Our resting state dataset does not include enough scanning sessions to concatenate and guarantee a larger number of datapoints than variables. To circumvent this dimensionality problem and be able to invert the covariance matrices, we applied FSL’s subsamp2 tool to downsample the MNI-registered functional data from 2mm isotropic voxels to 4mm isotropic voxels. The downsampling implied an eight-fold decrease in the number of voxels in the ROIs, thus reducing the number of variables in the covariance matrices to approximate 3,100 variables. This reduction was sufficient to obtain enough concatenated datapoints for the voxelwise conditional independence tests. The enlarged voxels may also aid with variations across individuals in the roles of neighboring voxels and with small co-registration errors. Results for left hemisphere high communication sub-regions in the caudate and in the putamen with OFC, DLPFC and posterior parietal cortex are shown in figures 14 - 16. Movies 6 – 11 show corresponding rotating views. Results for right hemisphere are given in figures S9 – S11 in supplementary material.

**Figure 14.**
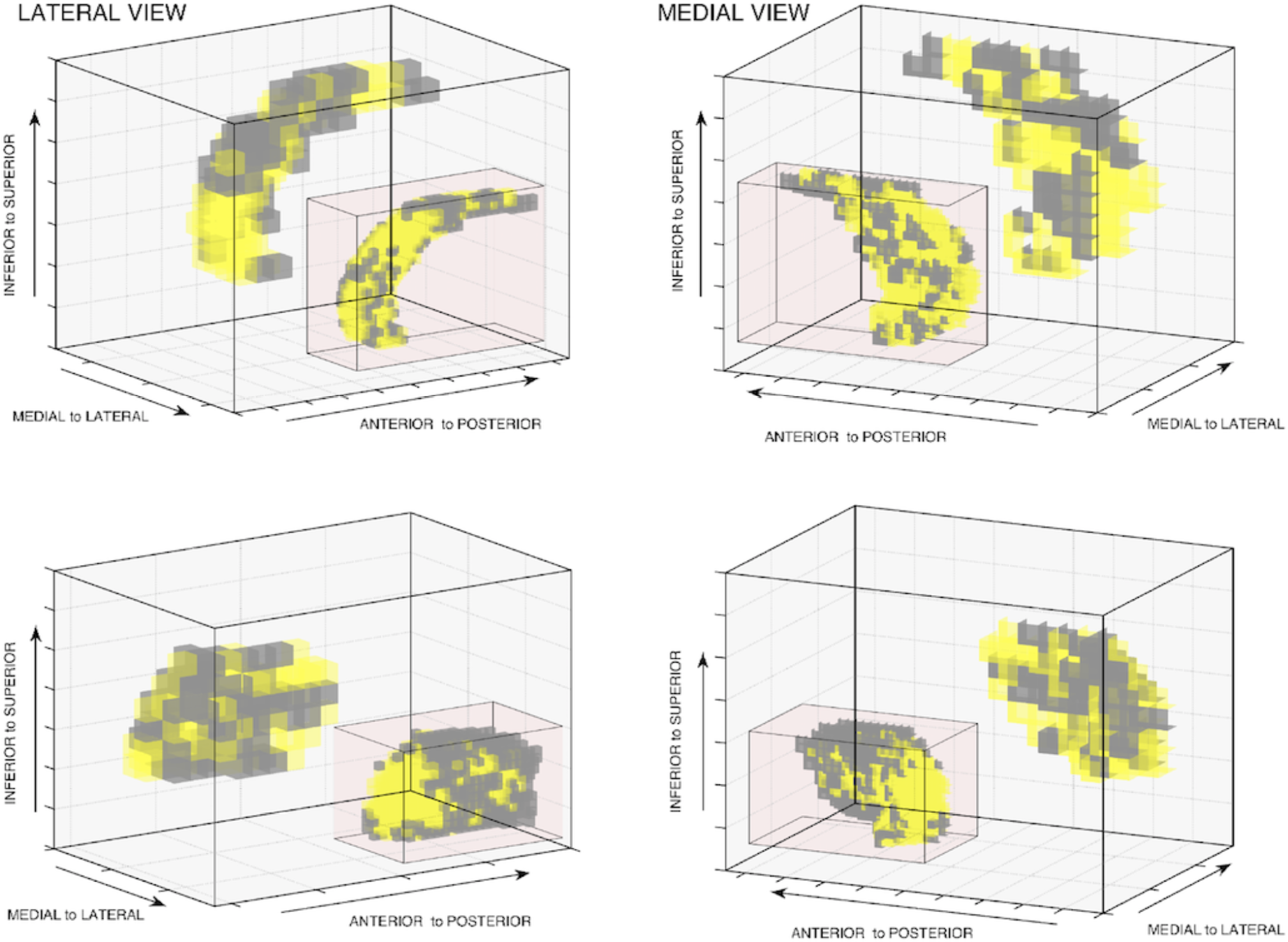
Main boxes show 3D voxel space representations of left hemisphere functional high communication sub-regions (yellow) in the caudate (top row) and in the putamen (bottom row), for orbitofrontal cortex (OFC) at 4mm voxel resolution. Inset boxes show the corresponding structural connectivity endpoints with OFC obtained by Jarbo and Verstynen (2015) at 2mm voxel resolution. See movie 6 and 7.

**Figure 15.**
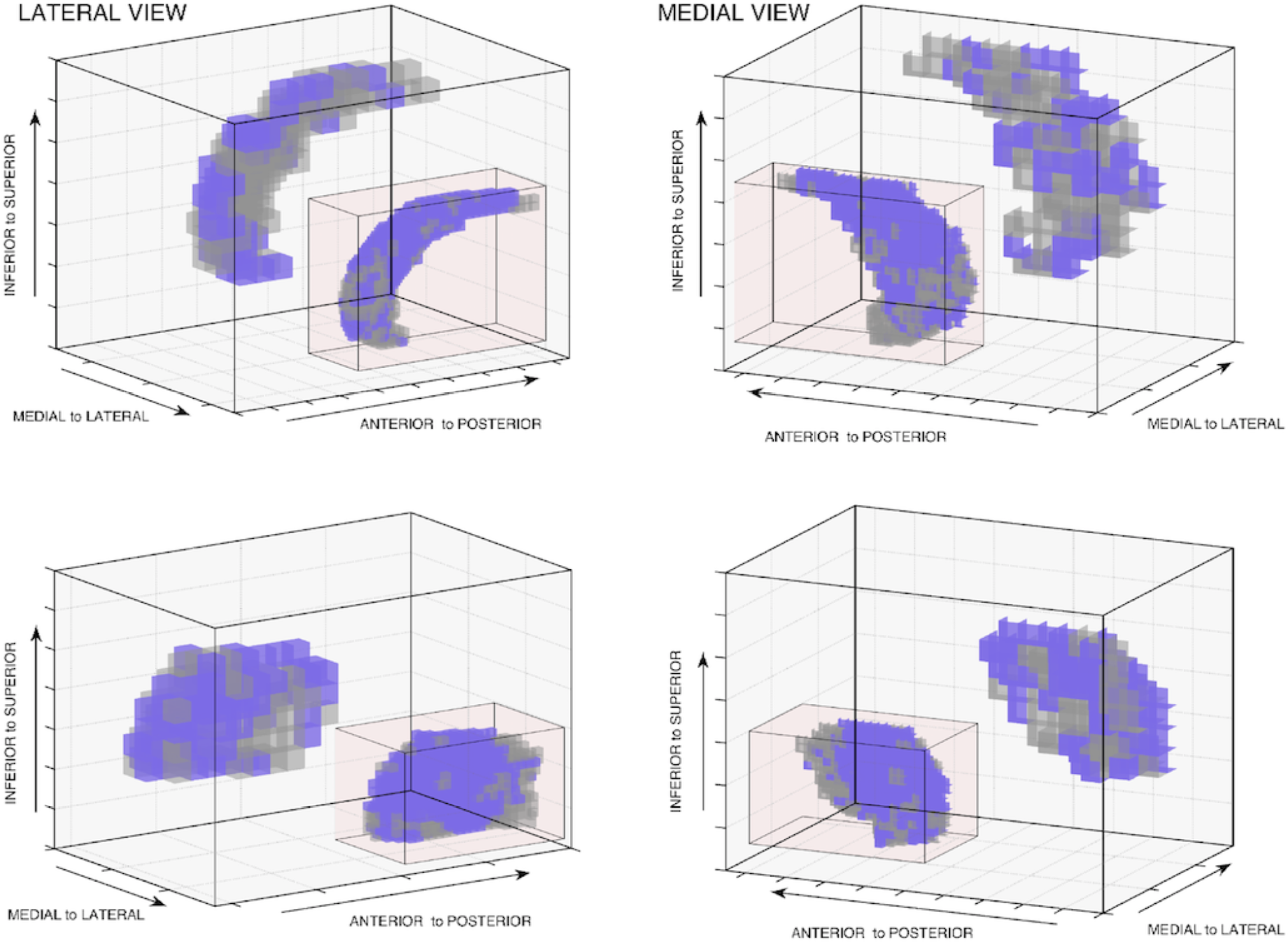
Main boxes show 3D voxel space representation of left hemisphere functional high communication sub-regions (purple) in the caudate (top row) and in the putamen (bottom row), for dorsolateral prefrontal cortex (DLPFC) at 4mm voxel resolution. Inset boxes show the corresponding structural connectivity endpoints with DLPFC obtained by Jarbo and Verstynen (2015) at 2mm voxel resolution. See movie 8 and 9.

**Figure 16.**
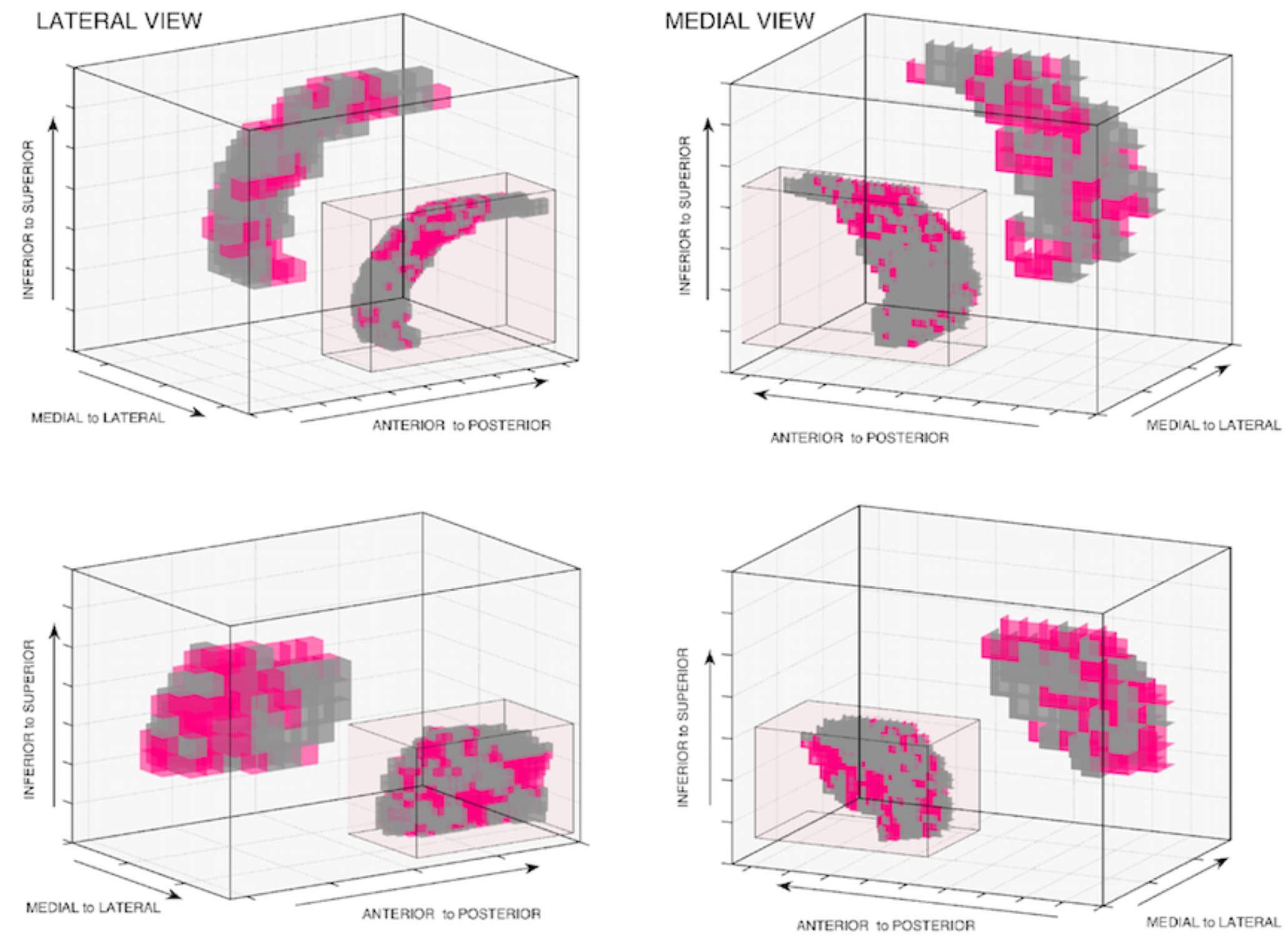
Main boxes show 3D voxel space representation of left hemisphere functional high communication sub-regions (pink) in caudate (top row) and in putamen (bottom row), for posterior parietal cortex (parietal) at 4mm voxel resolution. Inset boxes show corresponding structural connectivity endpoints with parietal cortex obtained by Jarbo and Verstynen (2015) at 2mm voxel resolution. See movie 10 and 11.

To contrast our functional results to the structural results of Jarbo and Verstynen (2015), we use NIfTI images at MNI 2mm space for group corticostriatal structural connectivity endpoints (see Jarbo and Verstynen (2015) for details on their group structural analysis), and mask these images with the AAL ROIs for caudate and putamen to extract voxelwise structural connectivity endpoints for each of the OFC, DLPFC and parietal cases. Caudate and putamen ROIs are represented in 3D voxel space plots at their original 2mm voxel resolution, showing in an assigned color, voxels with cortical connectivity endpoints for the OFC, DLPFC and parietal cases. For comparison, each of these voxelwise structural connectivity endpoints 3D plots was included as an inset box in the corresponding caudate and putamen functional connectivity 3D plot in figures 14 – 16.

## Discussion

In the hippocampus network case, concatenating ten medial temporal lobe resting state high resolution scans of the same individual, our Voxelwise Conditional Independence algorithm finds on purely statistical grounds, for each pair of regions of interest, spatially proximate voxels in different functional regions that are highly connected to neighboring and distant voxels. Qualitatively similar results are obtained with as few as five scans (see figure S4 and S8 in supplementary material). These results are in good agreement with those suggested by high resolution diffusion tensor imaging (Zeineh et al., 2012), and they are qualitatively stable under misalignments of voxels by 1 or 2 voxel position in any direction. In the corticostriatal network case, downsampling the size of the voxels and concatenating resting state data from multiple individuals, we are able to recover voxel to voxel functional connections in the caudate and the putamen that follow qualitatively voxelwise patterns of structural connectivity from deterministic fiber tractography on diffusion spectrum imaging. The data analysis procedures of course differ and this may explain some of the discrepancies observed. The group structural connectivity conclusions in Jarbo and Verstynen (2015) were obtained by t-tests over 59 individuals, whereas we have concatenated BOLD time series data from 50 of those individuals. The downsampling of the voxel size in our functional analysis and small spatial differences in the regions of interest possibly introduces some variations. In the illustrations above, we thresholded our results to select voxels with high functional connectivity degree, and while many of the highly functionally connected voxels correspond to structurally connected voxels, it may be that some of the structurally connected voxels have lower functional connectivity degree and thus were not included in the high communication sub-region. We also notice that in some cases voxels included in high communication sub-regions seem to lack structural connections, particularly in the posterior area of the putamen and in the utmost inferior area of the caudate. One plausible explanation is the presence of latent confounding variables that produce spurious functional connection estimates because they are not properly conditioned on. This scenario would arise if, for example, a cortical region of interest with inputs to the putamen and connected with the orbitofrontal cortex and with the dorsolateral prefrontal cortex is not properly included in our corticostriatal causal graph.

Our procedure can be thought as an application of partial correlation estimation of functional connectivity (Salvador et al., 2005; Marrelec et al., 2006; Liu et al., 2008; Ferrarini et al., 2009; Marrelec and Fransson, 2011; Smith et al., 2011, 2013; Brier et al., 2015) but at voxel level scale, with a restricted set of variables in the conditioning set derived from a given ROI level causal graph and with the goal of detecting subsets of highly connected voxels within pairs of assumed causally directly connected ROIs.

The analyses of BOLD data described here illustrate one method for resolving interneural processes at the voxel level, that is, establishing causal connections between individual voxels. We have used the technique to estimate high communication sub-regions based on the voxelwise connectivity degree, but it could be used for many other purposes, for example for identifying the internal causal structure of regions of interest or for identifying voxels near the physical boundaries of a region of interest that ought to be included within it or excluded from it.

Alternative methods for the same purposes are possible and deserve to be investigated. Other conditional independence procedures with our strategy might use penalized sparse partial regression methods such as adaptive lasso (Zou, 2006) or glasso (Friedman et al., 2008), and recent implementations such as QUIC and BigQUIC allow to solve problems for up to one million variables (Hsieh et al., 2013, 2014). These methods have the disadvantage that a sparsity parameter needs to be chosen independently for each individual and pair of regions considered; and cross-validated parameter selection require extensive computation for sets of variables of the size inspected here. The conditional independence method we have used has the advantages that it is reasonably fast and can be used with a false discovery rate for statistical error control, but it has some disadvantages. It requires a sample size large enough to avoid dimensionality restrictions, which, as in our applications, may require concatenating scans. Concatenation itself can produce non-causal associations (Ramsey, et al., 2011). This problem may be reduced with high-dimensional methods such as glasso. Further, all conditional independence methods require prior knowledge of the directed graph of effective connections between regions of interest to properly define the dependencies that should be tested. Without such knowledge, partial correlation analysis introduces false positive connections between pairs of variables that have a common effect (Mumford and Ramsey, 2014). The methods therefore should be used only with some care when estimating effective connections at the voxel level. These limitations can in principle be addressed with the IMaGES algorithm (Ramsey, et al 2010, 2011), which, however, does not produce error probabilities, or with the PC algorithm (Spirtes and Glymour, 1991), or its modifications (Colombo and Maathuis, 2014), which allow false discovery rate error probabilities but are comparatively slow and require a method for conditional independence testing on multiple independent data sets (Fisher, 1950; Tillman, 2009). The computational challenges facing these several methods can in principle be addressed by parallelization on multiple-thread supercomputers (Ramsey, 2015).

In contrast to conditional independence procedures, simple correlation methods are expected to produce misleadingly dense connections because they confound multiple pathways between and within multiple regions of interest. For example, correlation does not identify the direct pathways from, e.g., dorsolateral prefrontal cortex to caudate, and the functional connectivity spatial patterns do not accord with the structural results. Even with a very low alpha level of 10^−25^ for the correlation significance test, the putamen and caudate connectivity pattern for each of the cortical inputs form a dense cluster of high-degree voxels located in the exactly same spatial region. In the medial temporal lobe data, using correlations with a very low alpha level of 10^−14^ for the correlation significance tests produces voxels with very high connectivity degree distributed across large areas of the corresponding ROIs without the tight spatial coherency observed in the high communication sub-regions obtained by our algorithm. This is probably an artifact of the transitive closure of correlations among variables in causal chains and from correlation of variables sharing a common cause.

## Conclusion

Properly selected regions of interest contain voxels of neural cells that act more or less coherently to affect neural cells in voxels in other regions of interest, but the working of sub-groups of neural cells internal to a ROI is not uniform in its effects on cells in other ROIs. Some collections of neural cells will more directly receive incoming signals from an external source; some will send to an external source or sources; some will chiefly pass signals within a ROI; and some will have multiple roles. The specialization is not all-or-none but a matter of degree. fMRI regions of interest decomposition with multiple scans can to some degree distinguish these sub-regions and bring us closer to understanding the functional and causal roles of neural complexes.

## Footnotes

1 The proper clustering of voxels into regions of interest from BOLD signals has been controversial, chiefly because of concerns about multiple hypothesis testing (Bennett et al., 2009; Zhang, 2013; Friston et al., 2002; Poldrack, 2007; Etzel et al., 2009). There are, however standard remedies, either in statistical software (e.g., Jenkinson et al., 2012; Penny et al., 2011) or by use of the False Discovery Rate (Benjamini and Hochberg, 1995) in combination with very small alpha values.

2 If two voxels *x* and *y* are not dependent and they have a common effect z, conditioning on voxel *z* will create a spurious association between *x* and *y*.

3 The original fMRI resting-state data of Jarbo and Verstynen (2015) contained 55 individuals. We removed an additional 5 individuals because of difficulties in the MNI standard space registration produced by non-brain structures.

## Acknowledgments

Research for this paper was supported by a grant from the James S. McDonnell Foundation and by the National Institute of Health under Award Number U54HG008540 and Award Number 1R01LM012087-01. The content is solely the responsibility of the authors and does not necessarily represent the official views of the National Institutes of Health. We thank Russell Poldrack for sharing of the medial temporal lobe data and helpful advice; and Timothy Verstynen and Patrick Beukema from the Cognitive Axon Lab at Carnegie Mellon University, for sharing of the corticostriatal data and helpful advice.

## Author contributions

R.S.R, J.D.R and C.G conceived the algorithm, analyzed the data and wrote the paper; J.C.L defined the human medial temporal lobe regions of interest. K.J. provided the raw resting state fMRI and structural connectivity endpoints data for the corticostriatal case. R.S.R. prepare all the figures.

## APPENDIX

Technical Remarks on the Voxelwise Conditional Independence Algorithm.

(a) In the medial temporal lobe case, the α level used for the False Discovery Rate procedure in step 7 of the algorithm is quite low. The reason is that for our medial temporal lobe resting state fMRI data, distributions of the BOLD signals of individual voxels are moderately to extremely non-Gaussian according to the Anderson-Darling test of non-Gaussianity. As a result, the Gaussian Fisher Z transform does not apply; however, a generalization of the Fisher Z transform does apply (Hawkins et al., 1989).

Computation of the Generalized Fisher Z is much more time consuming than computation of the Gaussian Fisher Z. Instead, conditional independence relations for all voxels in two regions of interest were computed for both Gaussian Fisher Z and Generalized Fisher Z, finding that a Generalized Fisher Z cutoff index *k* of the FDR procedure at an α level of 0.001corresponded to Gaussian Fisher Z p-values in the range of 10^−14^ - 10^−12^. Thus, using a Gaussian Fisher Z, we choose an α level for FDR of 10^−14^ for the medial temporal lobe data in all cases. Choice of a higher α level for the Generalized Fisher Z comparisons would of course yield higher values for the Gaussian Fisher Z, with correspondingly higher connectivity degrees.

(b) The Gaussian CDF in principle never has the value 1, but all numerical computational implementations return 1 for extreme positive values of the statistic. Values of 1 for the CDF correspond to p-values of 0, and hence computation of p-values does not distinguish among very low values. Therefore, at a small risk of false positives (incorrect rejections of the conditional independence hypothesis) we ignore the possibility that in the FDR procedure, real values of computed p-values of 0 could be > kα/m, where k is the index of the hypothesis test in the sorted list of p-values and m is the number of tests.

(c) In step 4 of the algorithm, the discounting of sample size in the calculation of the Gaussian Fisher Z for the partial correlations follows Spirtes et al., (2000).

(d) Note that r(*x, y*) in step 3 of the algorithm is the correlation of *x* and *y* conditional on V \ <*x, y*>; where V is the set of all the voxels in R_x_, R_y_, and in the conditioning separating subset <R_z1_,…,R_z*n*_>; this conditional (or partial) correlation is easily calculated from the precision matrix, Q, and we use it here.

(e) In some cases, as with the corticostriatal data used here, the number of voxels to be compared by method (a) above, for the selection of α level for the FDR, is infeasibly large. Instead, we chose the *smallest α level that results in a non-singular distribution of degrees of connectivity of the voxels.* For the corticostriatal data we select an α level for FDR of 0.001.

(f) For the False Discovery Rate computation we follow the Benjamini-Hochberg procedure (Benjamini and Hochberg, 1995).

(g) The voxel connectivity degree empirical distribution can be very different among ROIs, varying in shape and domain depending on intrinsic connectivity properties of the regions of interest. For this reason, in step 9 of the algorithm we must compute the k-means clustering independently for each ROI connectivity degree distribution.

(h) To guarantee the robustness of the k-means clustering result for each case, we run the k-means algorithm 1,000 times and choose the mode.

(i) The algorithm was implemented in Java, and run in a 1.4Ghz dual core, 8G RAM, MacBook Air.

(j) Java code for the algorithm is currently available on request but will be placed in a GitHub repository.

(k) ROIs 3D renders and 3D voxel space figures were done in MATLAB R2015a, using functions *patch* and *isonormals*, and the package *vol3d v2.*

## References

Amaral, D. G. (1993). Emerging principles of intrinsic hippocampal organization. Current opinion in neurobiology, 3(2), 225–229.

Beckmann, C. F., & Smith, S. M. (2004). Probabilistic independent component analysis for functional magnetic resonance imaging. Medical Imaging, IEEE Transactions on, 23(2), 137–152.

Benjamini, Y., & Hochberg, Y. (1995). Controlling the false discovery rate: a practical and powerful approach to multiple testing. Journal of the Royal Statistical Society. Series B (Methodological), 289–300.

Bennett, C. M., Wolford, G. L., & Miller, M. B. (2009). The principled control of false positives in neuroimaging. Social cognitive and affective neuroscience, 4(4), 417–422.

Brier, M. R., Mitra, A., McCarthy, J. E., Ances, B. M., & Snyder, A. Z. (2015). Partial covariance based functional connectivity computation using Ledoit–Wolf covariance regularization. NeuroImage, 121, 29–38.

Canto, C. B., Koganezawa, N., Beed, P., Moser, E. I., & Witter, M. P. (2012). All layers of medial entorhinal cortex receive presubicular and parasubicular inputs. The Journal of Neuroscience, 32(49), 17620–17631.

Colombo, D., & Maathuis, M. H. (2014). Order-independent constraint-based causal structure learning. The Journal of Machine Learning Research, 15(1), 3741–3782.

Coulter, D. A., Yue, C., Ang, C. W., Weissinger, F., Goldberg, E., Hsu, F. C., … & Takano, H. (2011). Hippocampal microcircuit dynamics probed using optical imaging approaches. The Journal of physiology, 589(8), 1893–1903.

Ding, S. L., & Van Hoesen, G. W. (2010). Borders, extent, and topography of human perirhinal cortex as revealed using multiple modern neuroanatomical and pathological markers. Human brain mapping, 31(9), 1359–1379.

Ekstrom, A. D., Bazih, A. J., Suthana, N. A., Al-Hakim, R., Ogura, K., Zeineh, M., Burggren, A.C., & Bookheimer, S. Y. (2009). Advances in high-resolution imaging and computational unfolding of the human hippocampus. Neuroimage, 47(1), 42–49.

Etzel, J. A., Gazzola, V., & Keysers, C. (2009). An introduction to anatomical ROI-based fMRI classification analysis. Brain research, 1282, 114–125.

Faria, A. V., Joel, S. E., Zhang, Y., Oishi, K., van Zjil, P., Miller, M. I., Pekar, J.J., & Mori, S. (2012). Atlas-based analysis of resting-state functional connectivity: Evaluation for reproducibility and multi-modal anatomy–function correlation studies. Neuroimage, 61(3), 613–621.

Ferrarini, L., Veer, I. M., Baerends, E., van Tol, M. J., Renken, R. J., van der Wee, N. J., … & Milles, J. (2009). Hierarchical functional modularity in the resting-state human brain. Human brain mapping, 30(7), 2220–2231.

Fisher, R.A. (1950). Statistical methods for research workers. London: Oliver and Boyd. 11^th^ edition.

Friedman, J., Hastie, T., & Tibshirani, R. (2008). Sparse inverse covariance estimation with the graphical lasso. Biostatistics, 9(3), 432–441.

Friston, K. J., Glaser, D. E., Henson, R. N., Kiebel, S., Phillips, C., & Ashburner, J. (2002). Classical and Bayesian inference in neuroimaging: applications. Neuroimage, 16(2), 484–512.

Griffanti, L., Salimi-Khorshidi, G., Beckmann, C. F., Auerbach, E. J., Douaud, G., Sexton, C. E., … & Moeller, S. (2014). ICA-based artefact removal and accelerated fMRI acquisition for improved resting state network imaging. Neuroimage, 95, 232–247.

Hawkins, D. L. (1989). Using U statistics to derive the asymptotic distribution of Fisher’s Z statistic. The American Statistician, 43(4), 235–237.

Hsieh, C. J., Sustik, M. A., Dhillon, I. S., Ravikumar, P. K., & Poldrack, R. (2013). BIG & QUIC: Sparse inverse covariance estimation for a million variables. In Advances in Neural Information Processing Systems (pp. 3165–3173).

Hsieh, C. J., Sustik, M. A., Dhillon, I. S., & Ravikumar, P. (2014). QUIC: quadratic approximation for sparse inverse covariance estimation. The Journal of Machine Learning Research, 15(1), 2911–2947.

Insausti, R., & Amaral, D. G. (2004) Hippocampal formation. In Paxinos, G. & Mai, J.K., (eds.), The Human Nervous System, 2^nd^ Edition, Academic Press.

Jarbo, K., & Verstynen, T. D. (2015). Converging structural and functional connectivity of orbitofrontal, dorsolateral prefrontal, and posterior parietal cortex in the human striatum. The Journal of Neuroscience, 35(9), 3865–3878.

Jenkinson, M., & Smith, S. (2001). A global optimisation method for robust affine registration of brain images. Medical image analysis, 5(2), 143–156.

Jenkinson, M., Bannister, P., Brady, M., & Smith, S. (2002). Improved optimization for the robust and accurate linear registration and motion correction of brain images. Neuroimage, 17(2), 825–841.

Jenkinson, M., Beckmann, C. F., Behrens, T. E., Woolrich, M. W., & Smith, S. M. (2012). FSL. Neuroimage, 62(2), 782–790.

Kerr, K. M., Agster, K. L., Furtak, S. C., & Burwell, R. D. (2007). Functional neuroanatomy of the parahippocampal region: the lateral and medial entorhinal areas. Hippocampus, 17(9), 697–708.

Laumann, T. O., Gordon, E. M., Adeyemo, B., Snyder, A. Z., Joo, S. J., Chen, M. Y., … & Schlaggar, B. L. (2015). Functional system and areal organization of a highly sampled individual human brain. Neuron, 87(3), 657–670.

Liang, J. C., Wagner, A. D., & Preston, A. R. (2013). Content representation in the human medial temporal lobe. Cerebral Cortex, 23(1), 80–96.

Libby, L. A., Ekstrom, A. D., Ragland, J. D., & Ranganath, C. (2012). Differential connectivity of perirhinal and parahippocampal cortices within human hippocampal subregions revealed by high-resolution functional imaging. The Journal of Neuroscience, 32(19), 6550–6560.

Liu, Y., Liang, M., Zhou, Y., He, Y., Hao, Y., Song, M., … & Jiang, T. (2008). Disrupted small-world networks in schizophrenia. Brain, 131(4), 945–961.

Malykhin, N. V., Lebel, R. M., Coupland, N. J., Wilman, A. H., & Carter, R. (2010). In vivo quantification of hippocampal subfields using 4.7 T fast spin echo imaging. Neuroimage, 49(2), 1224–1230.

Marrelec, G., Krainik, A., Duffau, H., Pélégrini-Issac, M., Lehéricy, S., Doyon, J., & Benali, H. (2006). Partial correlation for functional brain interactivity investigation in functional MRI. Neuroimage, 32(1), 228–237.

Marrelec, G., & Fransson, P. (2011). Assessing the influence of different ROI selection strategies on functional connectivity analyses of fMRI data acquired during steady-state conditions. PLoS One, 6(4), e14788.

Moeller, S., Yacoub, E., Olman, C. A., Auerbach, E., Strupp, J., Harel, N., & Uğurbil, K. (2010). Multiband multislice GE-EPI at 7 tesla, with 16-fold acceleration using partial parallel imaging with application to high spatial and temporal whole-brain fMRI. Magnetic Resonance in Medicine, 63(5), 1144–1153.

Mumford, J. A., & Ramsey, J. D. (2014). Bayesian networks for fMRI: a primer. Neuroimage, 86, 573–582.

Nieto-Castanon, A., Ghosh, S. S., Tourville, J. A., & Guenther, F. H. (2003). Region of interest based analysis of functional imaging data. Neuroimage, 19(4), 1303–1316.

Penny, W. D., Friston, K. J., Ashburner, J. T., Kiebel, S. J., & Nichols, T. E. (Eds.). (2011). Statistical Parametric Mapping: The Analysis of Functional Brain Images. Academic Press.

Poldrack, R. A. (2007). Region of interest analysis for fMRI. Social cognitive and affective neuroscience, 2(1), 67–70.

Poldrack, R. A., Laumann, T. O., Koyejo, O., Gregory, B., Hover, A., Chen, M. Y., … & Hunicke-Smith, S. (2015). Long-term neural and physiological phenotyping of a single human. Nature communications, 6.

Preston, A. R., & Wagner, A. D. (2007). The medial temporal lobe and memory. In Martinez Jr, J.L., & Kesner, R. P. (eds.), Neurobiology of learning and memory, 2^nd^ Edition, Academic Press.

Preston, A. R., Bornstein, A. M., Hutchinson, J. B., Gaare, M. E., Glover, G. H., & Wagner, A. D. (2010). High-resolution fMRI of content-sensitive subsequent memory responses in human medial temporal lobe. Journal of Cognitive Neuroscience, 22(1), 156–173.

Ramsey, J. D., Hanson, S. J., Hanson, C., Halchenko, Y. O., Poldrack, R. A., & Glymour, C. (2010). Six problems for causal inference from fMRI. Neuroimage, 49(2), 1545–1558.

Ramsey, J. D., Hanson, S. J., & Glymour, C. (2011). Multi-subject search correctly identifies causal connections and most causal directions in the DCM models of the Smith et al. simulation study. NeuroImage, 58(3), 838–848.

Ramsey, J. D., Spirtes, P., & Glymour, C. (2011). On meta-analyses of imaging data and the mixture of records. NeuroImage, 57(2), 323–330.

Ramsey, J. D. (2015). Scaling up Greedy Causal Search for Continuous Variables. arXiv preprint arXiv:1507.07749.

Rohlfing, T., Zahr, N. M., Sullivan, E. V., & Pfefferbaum, A. (2010). The SRI24 multichannel atlas of normal adult human brain structure. Human brain mapping, 31(5), 798–819.

Salimi-Khorshidi, G., Douaud, G., Beckmann, C. F., Glasser, M. F., Griffanti, L., & Smith, S. M. (2014). Automatic denoising of functional MRI data: combining independent component analysis and hierarchical fusion of classifiers. Neuroimage, 90, 449–468.

Salvador, R., Suckling, J., Coleman, M. R., Pickard, J. D., Menon, D., & Bullmore, E. D. (2005). Neurophysiological architecture of functional magnetic resonance images of human brain. Cerebral cortex, 15(9), 1332–1342.

Smith, S. M. (2002). Fast robust automated brain extraction. Human brain mapping, 17(3), 143–155.

Smith, S. M., Miller, K. L., Salimi-Khorshidi, G., Webster, M., Beckmann, C. F., Nichols, T. E., Ramsey, J. D., & Woolrich, M. W., (2011). Network modelling methods for FMRI. Neuroimage, 54(2), 875–891.

Smith, S. M., Vidaurre, D., Beckmann, C. F., Glasser, M. F., Jenkinson, M., Miller, K. L., Nicholson, T. E., & Van Essen, D. C. (2013). Functional connectomics from resting-state fMRI. Trends in cognitive sciences, 17(12), 666–682.

Spirtes, P., & Glymour, C. (1991). An algorithm for fast recovery of sparse causal graphs. Social science computer review, 9(1), 62–72.

Spirtes, P. Glymour, C., and Scheines, R., (2000). Causation, Prediction, and Search. MIT Press.

Squire, L. R., Stark, C. E., & Clark, R. E. (2004). The medial temporal lobe. Annu. Rev. Neurosci., 27, 279–306.

Tzourio-Mazoyer, N., Landeau, B., Papathanassiou, D., Crivello, F., Etard, O., Delcroix, N., … & Joliot, M. (2002). Automated anatomical labeling of activations in SPM using a macroscopic anatomical parcellation of the MNI MRI single-subject brain. Neuroimage, 15(1), 273–289.

Tillman, R. E. (2009). Structure learning with independent non-identically distributed data. In Proceedings of the 26th Annual International Conference on Machine Learning (pp. 1041–1048). ACM.

Van Essen, D. C., Ugurbil, K., Auerbach, E., Barch, D., Behrens, T. E. J., Bucholz, R., … & Yacoub, E. (2012). The Human Connectome Project: a data acquisition perspective. Neuroimage, 62(4), 2222–2231.

Wilson, C. J. (1989). Basal Ganglia. In Shepherd, G. M. (ed.), The Synaptic Organization of the Brain, 4^th^ Edition, Oxford University Press.

Witter, M. P. (1993). Organization of the entorhinal — hippocampal system: A review of current anatomical data. Hippocampus, 3(S1), 33–44.

Witter, M. P., Naber, P. A., van Haeften, T., Machielsen, W. C., Rombouts, S. A., Barkhof, F., … & Lopes da Silva, F. H. (2000). Cortico-hippocampal communication by way of parallel parahippocampal-subicular pathways. Hippocampus, 10(4), 398–410.

Wong, E. C. (2014). Direct Imaging of Functional Networks. Brain connectivity, 4(7), 481–486.

Yassa, M. A., Stark, S. M., Bakker, A., Albert, M. S., Gallagher, M., & Stark, C. E. (2010). High-resolution structural and functional MRI of hippocampal CA3 and dentate gyrus in patients with amnestic Mild Cognitive Impairment. Neuroimage, 51 (3), 1242–1252.

Yushkevich, P. A., Avants, B. B., Pluta, J., Das, S., Minkoff, D., Mechanic-Hamilton, D., … & Detre, J. A. (2009). A high-resolution computational atlas of the human hippocampus from postmortem magnetic resonance imaging at 9.4 T. Neuroimage, 44(2), 385–398.

Yushkevich, P. A., Wang, H., Pluta, J., Das, S. R., Craige, C., Avants, B. B., … & Mueller, S. (2010). Nearly automatic segmentation of hippocampal subfields in *in vivo* focal T2-weighted MRI. Neuroimage, 53(4), 1208–1224.

Yushkevich, P. A., Amaral, R. S., Augustinack, J. C., Bender, A. R., Bernstein, J. D., Boccardi, M., … & Chételat, G. (2015). Quantitative comparison of 21 protocols for labeling hippocampal subfields and parahippocampal subregions in in vivo MRI: Towards a harmonized segmentation protocol. NeuroImage, 111, 526–541.

Zeineh, M. M., Engel, S. A., & Bookheimer, S. Y. (2000). Application of cortical unfolding techniques to functional MRI of the human hippocampal region. Neuroimage, 11 (6), 668–683.

Zeineh, M. M., Engel, S. A., Thompson, P. M., & Bookheimer, S. Y. (2001). Unfolding the human hippocampus with high resolution structural and functional MRI. The Anatomical Record, 265(2), 111–120.

Zeineh, M. M., Holdsworth, S., Skare, S., Atlas, S. W., & Bammer, R. (2012). Ultrahigh resolution diffusion tensor imaging of the microscopic pathways of the medial temporal lobe. Neuroimage, 62(3), 2065–2082.

Zhang, C. (2013). Multiple Comparison with Applications in Neuroimaging Data. Presentation at SAMSI 2013: NeuroImaging Data Analysis.

Zou, H. (2006). The adaptive lasso and its oracle properties. Journal of the American statistical association, 101 (476), 1418–1429.

